# Extracellular water withdrawal drives disease resistance in the phyllosphere

**DOI:** 10.64898/2026.08.27.747301

**Authors:** Charles Roussin-Léveillée, Sabrina Gauthier, Yezhou Hu, Jie Zhu, Faye Gaudreault- Lafleur, Soline Marty, Antoine Pelletier, Alexis Roy, Laurent D. Noël, Gitta L. Coaker, Xiufang Xin, Peter Moffett

## Abstract

A central question in immunity is how hosts arrest pathogen growth. Diverse plant pathogens create a water-soaked niche in host tissue essential for pathogenesis, yet how water shapes infection outcome is unknown. Using genetics and hyperspectral imaging, we show that extracellular water status is rate-limiting for both compatible and incompatible interactions. We find that the hypersensitive response of effector-triggered immunity (ETI) is, mechanistically, a desiccation event. Water loss imposes osmotic stress that arrests bacterial division while the pathogen remains alive and metabolically active, rather than killing it. Restoring apoplastic water reverses this stasis and licenses growth despite intact immune signaling and cell death. Water status, not immune signaling *per se*, gates pathogen growth. This reframes ETI as a controlled desiccation mechanism and identifies hydration as a decisive lever on disease outcome.

## INTRODUCTION

Manipulation of host immune and metabolic programmes by secreted effector proteins is essential for microbial pathogens of both animals and plants (*1, 2*). In plants, the induction of an effector-driven extracellular niche (EDEN), which consists of the enrichment of the extracellular space (i.e. the apoplast) in nutrients and water, is a determinant factor for growth of many pathogens (*3*). Water-rich extracellular niches induced by pathogens are commonly referred to as water-soaked lesions, which are visible to the naked eye. Water-soaking induction has been found to be triggered via diverse mechanisms, including stomatal manipulation via abscisic acid (ABA) signaling, delivery of water/nutrient channels to host cells, manipulation of nutrient export and cell-wall degradation (*4-11*). Notably, the effector family of protein AvrE, conserved from bacteria to certain oomycetes, has been identified alongside other effectors as a critical contributor to water soaking (*12, 13*).

Despite their importance in pathogenesis, it is still unclear how water-soaked lesions influence bacterial behavior and plant immune responses. To shed light on the function of water-soaking in plant-pathogen interactions, we combined hyperspectral imaging and conventional molecular biology approaches to address this longstanding question. This study reveals that the extracellular water status is a rate-limiting step for effective execution, but not activation of, plant immunity. We establish a new framework in which we propose that desiccation of plant tissues via intracellular immune receptor activation induces a water deprivation response in microbial pathogens forcing them to enter dormancy as a survival mechanism.

## Results

### Apoplastic water affects pattern-triggered immunity

Previous studies have shown that artificially maintaining a water-rich apoplast promotes bacterial growth and can compensate for the loss of certain effector proteins (*4, 13*). To test if apoplastic water might also affect plant defense responses, we evaluated the effect of maintaining the apoplast in a water-soaked (flooded) state on several immune outputs (Fig. 1A). Pretreatment of Arabidopsis leaves with the immunogenic peptide flg22 results in protection against a subsequent infection with *Pseudomonas syringae pv. tomato* (*Pst*) DC3000. However, this protection is entirely lost if the apoplastic state is maintained flooded immediately and continuously upon inoculation (Fig. 1, B and C). Maintaining the apoplast hydrated is not sufficient to restore full virulence to *Pst hrcC*^-^, a strain unable to deliver effectors (Fig 1D). This indicates that apoplastic water can abrogate the effects of pattern-triggered immunity (PTI) on a virulent strain of bacteria, but that effectors are required for additional growth-promoting purposes (*3, 14*). Maintaining the apoplast flooded did not prevent flg22 from triggering early immune responses as flg22 still elicited MAPK phosphorylation (Fig. 1E) and apoplastic ROS bursts, albeit reduced (Fig. 1, F and G), as well as apoplastic peroxidases (Fig. 1, H). Transcriptome analysis of Arabidopsis plants infected with WT *Pst* maintained in a flooded vs non-flooded state revealed that creating an artificially flooded environment does not induce a transcriptional signature that is distinct from naturally occurring water-soaking lesions (Fig. S1). This indicates that artificial flooding maintenance can be used as a water-soaking mimic.

**Figure 1:**
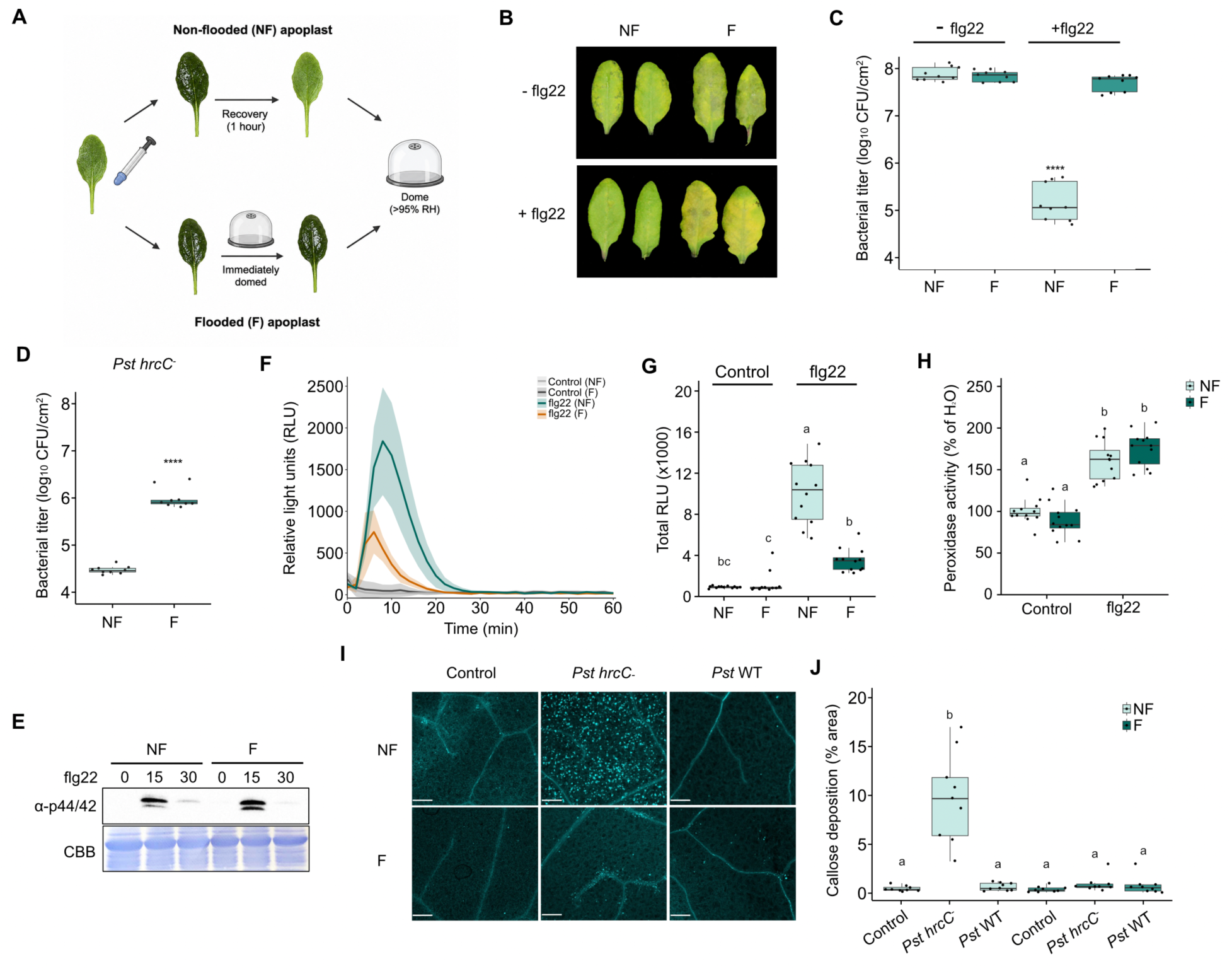
Apoplastic water affects pattern-triggered immune functions. **(A**) Model representing the methodology used to investigate the impact of apoplastic water on PTI. Briefly, Arabidopsis leaves were infiltrated with 10 mM MgCl2 (control) and was allowed to evaporate naturally for 1 hour to recover to the pre-infiltration non-flooded (NF) state, followed by covering with a plastic dome to maintain high humidity, or were immediately covered to avoid evaporation and maintain a flooded (F) state. (**B-C**) Apoplast flooding breaks flg22-mediated protection. Arabidopsis leaves were pre-infiltrated with water (-flg22) or with 1 μM flg22 and immediately allowed to dry. One day later, the same leaves were infiltrated with *Pst* WT (1 x 10^5^ CFU/mL) and maintained as NF or F with the *Pst* inoculum. Photos were taken (**B**) and bacteria counted (**C**) at 3 days post-infection (dpi). Asterisks indicate statistical significance (**** = p <0.0005) after performing a one-way ANOVA followed by Tukey’s HSD post-hoc test. (**D**) Apoplast flooding slightly enhances growth of a non-virulent bacterium. Bacterial count at 3 dpi of Arabidopsis leaves that were infiltrated with *Pst hrcC*^-^ at 1 x 10^5^ CFU/mL and apoplasts were maintained NF or F for 3 days. Asterisks indicate statistical significance (**** = p <0.0005) after performing a Student’s t-test. (**E**) Apoplast flooding does not alter MAPK phosphorylation post immune elicitation. Arabidopsis leaves were maintained in a NF or F state with 10 mM MgCl_2_ for 24 hours prior to being dried and treated with 1 μM flg22 for 0, 15 and 30 minutes. Total protein extracts were subsequently subjected to immunoblotting against MAPK p42/44. (**F-G**) Apoplast flooding negatively impacts immunity-induced ROS bursts. Arabidopsis leaf punches were maintained in a NF or F state for 24 hours prior to being treated with flg22 1 μM or ddH_2_O (control). ROS bursts were quantified over time (**F**) and as total relative light units (RLU) (**G**). Different letters indicate statistical significance (p < 0.05) after performing a Kruskal-Wallis followed by a Dunn’s post-hoc test (Bonferroni correction) statistical analysis. (**H**) Apoplast flooding does not impact extracellular peroxidase activity. Leaf discs from Arabidopsis leaf discs were pre-treated as in (F), followed by treatment with water (control) or 1 μM flg22 peptide. Total peroxidase activity was measured 20 h after treatment and data is shown as percentage of the water treated control (mean ± standard error [SE]). Different letters indicate statistical significance (p<0.05) after performing a one-way ANOVA followed by Tukey’s HSD post-hoc. (**I-J**) The apoplast water status influences callose deposition in response to bacteria. Confocal images (**I**) and count of callose as a percentage of deposit sites per acquired images (**J**) in Arabidopsis leaves were infiltrated with 10 mM MgCl_2_ (control) or with *Pst* WT or *hrcC*^-^ (1 x 10^8^ CFU/mL) and maintained in a NF or F state, as indicated. Inoculated areas were visualized by confocal microscopy (**I**) 3 days post-inoculation and callose was quantified as a percentage of deposit sites per acquired images (**J**). Scale bars represent 500 μm. Different letters indicate statistical significance (p < 0.05) after performing a Kruskal-Wallis followed by a Dunn’s post-hoc test (Bonferroni correction).

Callose deposition is a cell-wall based response induced by multiple biotic stresses, including bacteria defective in immunomodulatory effectors (*15*), such as *Pst hrcC*^-^. This response, however, is completely abrogated by *Pst* WT, or by maintaining a water flooded apoplastic space upon inoculation with *Pst hrcC*^-^ (Fig. 1I, J). This inhibition of callose deposition could be reversed upon exposure to ambient humidity levels, allowing apoplastic water evaporation, suggesting that the apoplast hydration status alone affects callose deposition (Fig. S2). This could be due to the dilution of key factors, however, maintaining the apoplast hydrated with hydrogen peroxide, Ca^2+^ or sorbitol did not rescue the production of callose. This suggests that a hydrated apoplast does not prevent callose production simply by diluting certain requisite molecules, or by affecting osmolarity (Fig. S3).

The water-soaking effectors HopM1 and AvrE1 are required to inhibit callose accumulation, as callose deposition was observed following infiltration with *Pst hopM1^-^/avrE1^-^*, similar to that seen with the *Pst hrcC^-^* mutant (Fig. S4). This is consistent with a previous report that the CEL cluster of effectors in *Pst*, which includes HopM1 and AvrE1, suppresses cell-wall based defenses in Arabidopsis (*15*). HopM1 and AvrE1 promote water-soaking by inducing ABA responses (*4, 5*) and these activities are essential for preventing callose deposition as *Pst* WT did not inhibit callose deposition in the Arabidopsis ABA biosynthetic mutant *aba2-1* (Fig. S5). The latter mutant shows constitutive callose accumulation and this, as well as *Pst*-induced callose, are inhibited simply by maintaining apoplast flooding, further inferring that apoplastic water alone is sufficient to prevent callose deposition (Fig. S5). These results indicate that apoplastic water does not directly affect immune signaling, but rather execution of plant defense, including cell-wall based reinforcement.

### Apoplastic water disrupts effective execution of ETI

Cell death initiated during the hypersensitive response (HR) has long been believed to be a major contributor to defense during effector-triggered immunity (ETI). This response is activated by NLR proteins and leads to pathogen growth arrest and disease resistance (*16, 17*). However, cell death is not always observed in ETI, particularly under conditions of low pathogen inoculum. Indeed, Arabidopsis leaves challenged with the avirulent strains *Pst* AvrRps4 and *Pst* AvrRpm1 displayed little to no visible signs of infection compared to leaves challenged with *Pst* WT (Fig. 2A). However, if apoplastic water was artificially maintained after pathogen challenge, leaves displayed strong disease symptoms whether they were infected with a virulent or avirulent *Pst* strain (Fig. 2A). Bacterial growth levels were found to be similar between *Pst* WT and *Pst* AvrRpm1 or AvrRps4 in flooded apoplast conditions, suggesting a complete break of NLR function (Fig. 2B). Similar results were observed in plants in which the apoplast flooding was induced by sealing the leaves with petroleum jelly, which leads to spontaneous water-soaking under high humidity by blocking transpiration, without requiring water injection (Fig. S6) (*18, 19*). To broaden our observations to another pathosystem, we evaluated the effect of water on avirulent *Xanthomonas campestris pv*. *campestris* CN06 WT strain (*20*). We found that leaf margin sealing and water-soaking significantly increased the virulence of the avirulent CN06 WT strain, as well as the virulent *Xanthomonas* CN06 *xopJ6* mutant, which lacks the effector XopJ6 recognized by an unknown resistance protein in cauliflower (Fig. S7).

**Figure 2:**
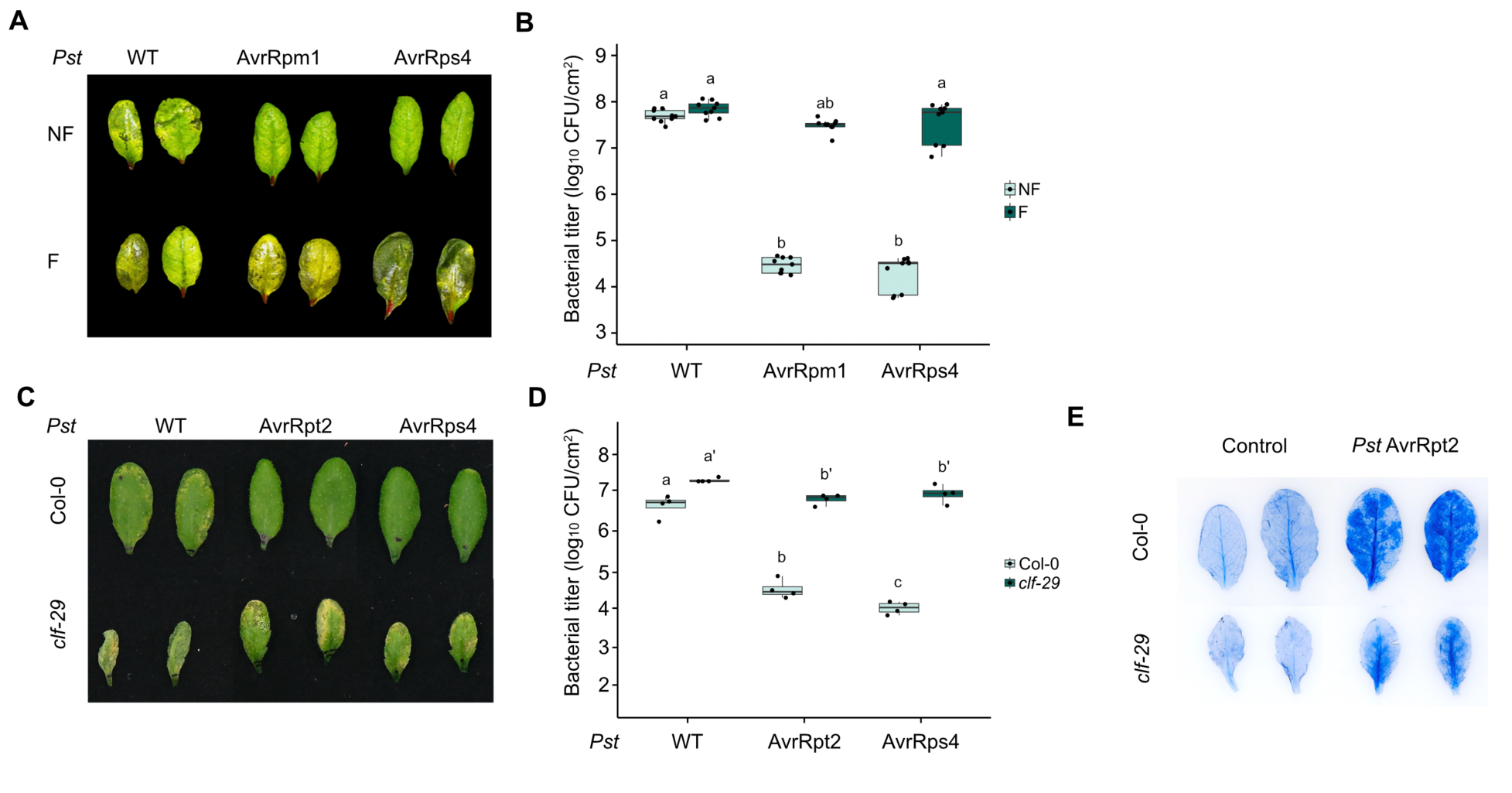
Apoplast hydration status uncouples ETI activation from execution. (**A-B**) Apoplast flooding breaks ETI-mediated disease prevention against *Pst*. Disease phenotype photos (**A**) and (**B**) bacterial count of Arabidopsis WT leaves that were infiltrated with *Pst* WT, *Pst* AvrRpm1 or *Pst* AvrRps4 (1 x 10^5^ CFU/mL) and maintained in a NF or F state for 3 days. Photos were taken and bacterial count evaluated at 3 dpi. Different letters indicate statistical significance (p < 0.05) after performing a Kruskal-Wallis followed by a Dunn’s post-hoc test (Bonferroni correction) test. (**C-D**) ETI-mediated protection against avirulent *Pst* strains is lost in the Arabidopsis spontaneous water-soaking mutant *clf*. Photos (**C**) and bacterial count (**D**) of Arabidopsis WT leaves that were infiltrated with *Pst* WT, *Pst* AvrRpt2 or *Pst* AvrRps4 (1 x 10^6^ CFU/mL) and maintained in a NF or F state for 3 days. Different letters indicate statistical significance (p < 0.05) after performing a one-way ANOVA followed by a Tukey’s HSD post-hoc test. (**E**) ETI-induced cell death in the Arabidopsis *clf-29* mutant. Leaves of four-week-old Arabidopsis WT and *clf-29* were infiltrated with *Pst* AvrRpt2 (1 x 10^8^ CFU/ml) or water (control) solution, plants were kept under high humidity (> 95% RH) after evaporation of excess water. Leaves were harvested and stained with trypan blue to observe dead cells. Photos were taken at 12 hours post infiltration.

To consolidate our findings on the impact of water on ETI, we studied ETI responses in the Arabidopsis *CURLYLEAF* (*clf*) mutant, which undergoes spontaneous water soaking at high humidity (*21*). Likewise, inoculation of *clf-29* with *Pst* WT, *Pst* AvrRpm1 and *Pst* AvrRps4 results in water soaking after 24 hours, whereas this is normally seen only in compatible interactions in WT Arabidopsis (Fig. S8A). Consistent with this, Arabidopsis *clf-29* mutants displayed clear disease phenotypes, as well as strong bacterial growth, three days after inoculation with *Pst* WT, *Pst* AvrRpm1 and *Pst* AvrRps4 strains (Fig. 2C, D). Interestingly, while ETI-associated cell death still occurred in the *clf-29* mutants (Fig. 2E), we observed a significant delay in tissue collapse (Fig. S8B, C). These observations suggest that extracellular water content reduces cell death symptoms (Fig. S8). The *clf-29* mutant has increased basal ABA biosynthesis levels (*21*). As such, we tested whether exogenous application may have the same effect. Exogenous application of ABA to the leaves of plants inoculated with *Pst* WT or AvrRpm1 revealed that ABA could partially rescue bacterial growth arrest during ETI (Fig. S9A). Supporting this observation, we found that *Pst* WT growth in the Arabidopsis *aba2-1* mutant was similar to *Pst* AvrRpm1 in WT plants (Fig. S9B), suggesting that a lack of virulence in both cases may be associated with an unfavourable apoplastic water status.

A role for ETI in blocking pathogen-induced water-soaking has been previously reported (*13, 22*). Indeed, this appears to be the case upon inoculation with low titers of *Pst* and *Xanthomonas* CN06, which represents more naturally occurring levels of bacteria at the start of the infection process (Fig. S7B and S10A). High titers of bacterial infiltration cause a breakdown of ETI, in that water-soaking-like lesions are observed, and bacterial growth reaches levels associated with a virulent infection (Fig. S10 A, B). These results serve as a cautionary note on evaluating ETI functions at high bacterial inoculum for studying physiologically relevant immune responses. Multiple mechanisms are likely at play in reducing water availability during ETI and PTI (*3*). To this effect, stomatal aperture analyses revealed that, in contrast to *Pst* WT, *Pst* AvrRpm1, AvrRps4 and AvrRpt2 do not induce stomatal closure at 24 hpi, even at high doses of inoculum (Fig. S10C). This observation begs the question of whether the water-soaking-like lesions observed with high dose of inoculum (OD_600_ = 0.02) at 24 hpi are indeed water-soaking. In almost every instance, we noticed the water-soaking-like ETI leaves to be ‘glossy’ at 24 hpi, similar to what is observed with *Pst* WT at a similar inoculum after 72 hpi. As this inoculum dosage leads to rapid tissue collapse after a humidity shift (*13*), we suggest that these are not true water-soaking lesions, but rather bursts of water leakage from dead plant cells. Together, these results indicate that although the HR occurs, ETI is functionally overwhelmed and does not initially limit pathogen growth due to the presence of this extracellular water.

### ETI-induced hypersensitive response leads to plant cell desiccation

High humidity has long been associated with a reduction in the hypersensitive response phenotype even when ETI limits microbial growth under such conditions (*23-25*). At the same time, ETI exists on a spectrum between HR and no visible HR depending on the magnitude of the ETI response and, presumably, the nature of the NLR/elicitor interaction (*26*). To better understand the role of water content in the control of infection during ETI, we performed hyperspectral imaging of leaves challenged with avirulent *Pst* strains carrying effectors inducing weak ETI responses (non-HR inducing; *Pst* HopAZ1 and *Pst* HopX1), as well as strains inducing a strong ETI response leading to a hypersensitive response (HR; *Pst* AvrRpm1 and *Pst* AvrRpt2) (*26*). After an incubation period of 16 hours under dome (>95% RH), leaf water content was monitored hourly for 12 hours following dome removal (Fig. 3A). We observed that tissue desiccation was strongly induced in plants inoculated with HR-eliciting strains, but not during infection with non-HR or WT strains (Fig. 3B, D and E). Nonetheless, non-HR-eliciting strains did not show water soaking. We also compared bacterial titers at 1 and 12 hours post-dome removal (humidity shift from >95% RH to 40% RH) and found that HR-eliciting strains exhibited growth stasis over this period (Fig. 3C). In contrast, non-HR strains did not show growth stasis for the same period, but still reached lower titers than the WT strain, indicating bacterial growth continued to be affected.

**Figure 3:**
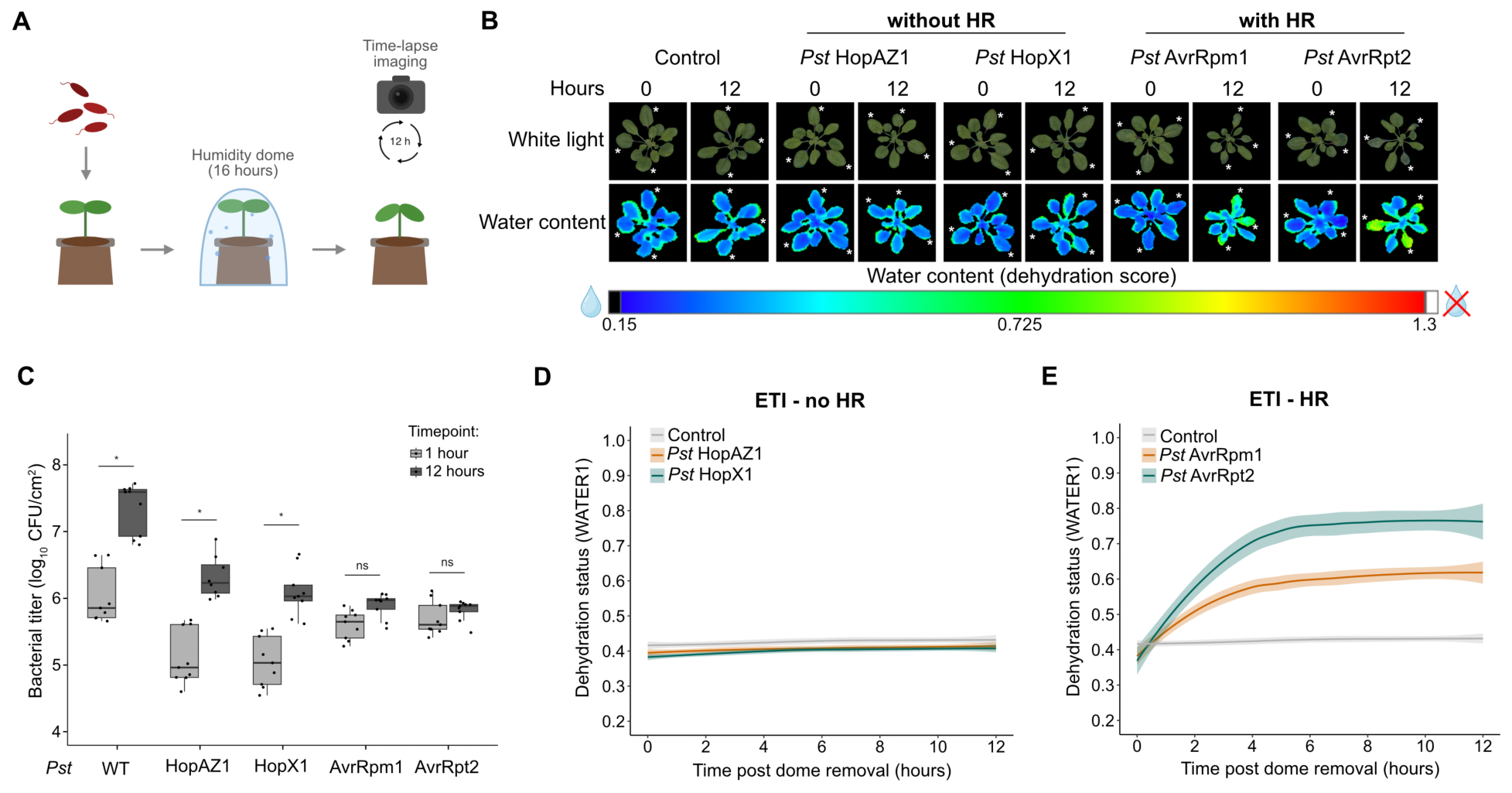
ETI-induced hypersensitive response leads to plant cell desiccation. (**A**) Model representing the methodology used to investigate the water content and hypersensitive response-driven desiccation during ETI. Briefly, *Arabidopsis* WT leaves were infiltrated with 10 mM MgCl_2_ (control), *Pst* HopAZ1, *Pst* HopX1, *Pst* AvrRpm1 or *Pst* AvrRpt2 (1 x 10^7^ CFU/ml) and kept at high humidity (>95% RH) under dome for 16 hours, then transferred to an automated phenotyping platform to acquire images every hour over a 12-hour period. (**B**) Hyperspectral and white light images of *Arabidopsis* WT plants as described in (**A**). WATER1 represents leaf water content (**C**) Bacterial titers at 1 and 12 hours post-dome removal from leaves as described in (**A-B**). Asterisks indicate statistical significance (p < 0.05) after performing a Mann-Whitney U test followed by a Bonferroni correction. (**D-E**) Quantification of the dehydration score of the leaves over time as observed in (**B**), of non-HR-eliciting strains (**D**) and HR-eliciting strains (**E**).

To specifically assess the HR-associated phenotype in the absence of bacterial infection, we performed hyperspectral imaging of plants expressing AvrRpm1 and AvrRps4 from dexamethasone– and estradiol-inducible promoters respectively. Following induction, plants were imaged hourly over a 25-hours period. We found that induction of either effector led to rapid and robust desiccation of plant tissues (Fig. S11). Together, these results indicate that the hypersensitive response associated with ETI leads to severe loss of water content in infected tissues, and that this desiccation correlates with bacterial growth arrest.

### ETI induces a reversible microbial growth stasis in the apoplast

It has previously been reported that a *Pst* WT strain in a resistant Arabidopsis ecotype, as well as non-pathogenic endophytes of the phyllosphere undergo microbial growth stasis under normal conditions (*27*). We evaluated whether *Pst* AvrRpm1 and *Pst* AvrRps4 strains would undergo a growth stasis over 10 days of infection rather than being eliminated by the plant immune system. We found that, while WT populations of *Pst* grew rapidly and could only be monitored until 3 dpi due to tissue maceration, avirulent and non-virulent *Pst* strains maintained population levels without growth over 10 days (Fig. S12). If extracellular water is rate-limiting to microbial growth in the apoplast, we posited that pathogenesis could be re-initiated by infiltrating water into the apoplast. We found that bacterial titers of *Pst* AvrRpm1 and AvrRps4 were maintained between 7 and 10 dpi, whether plants were kept under normal (70%) or high (>95%) relative humidity levels (Fig. 4A-C). However, a one-time pulse of infiltrating water into the apoplast was sufficient to re-activate bacterial growth to significant levels (near pathogenic), while maintaining water for 3 days in the apoplast post-infiltration at 7 dpi caused WT-level *Pst* growth and disease in Arabidopsis (Fig. 4A-C).

**Figure 4:**
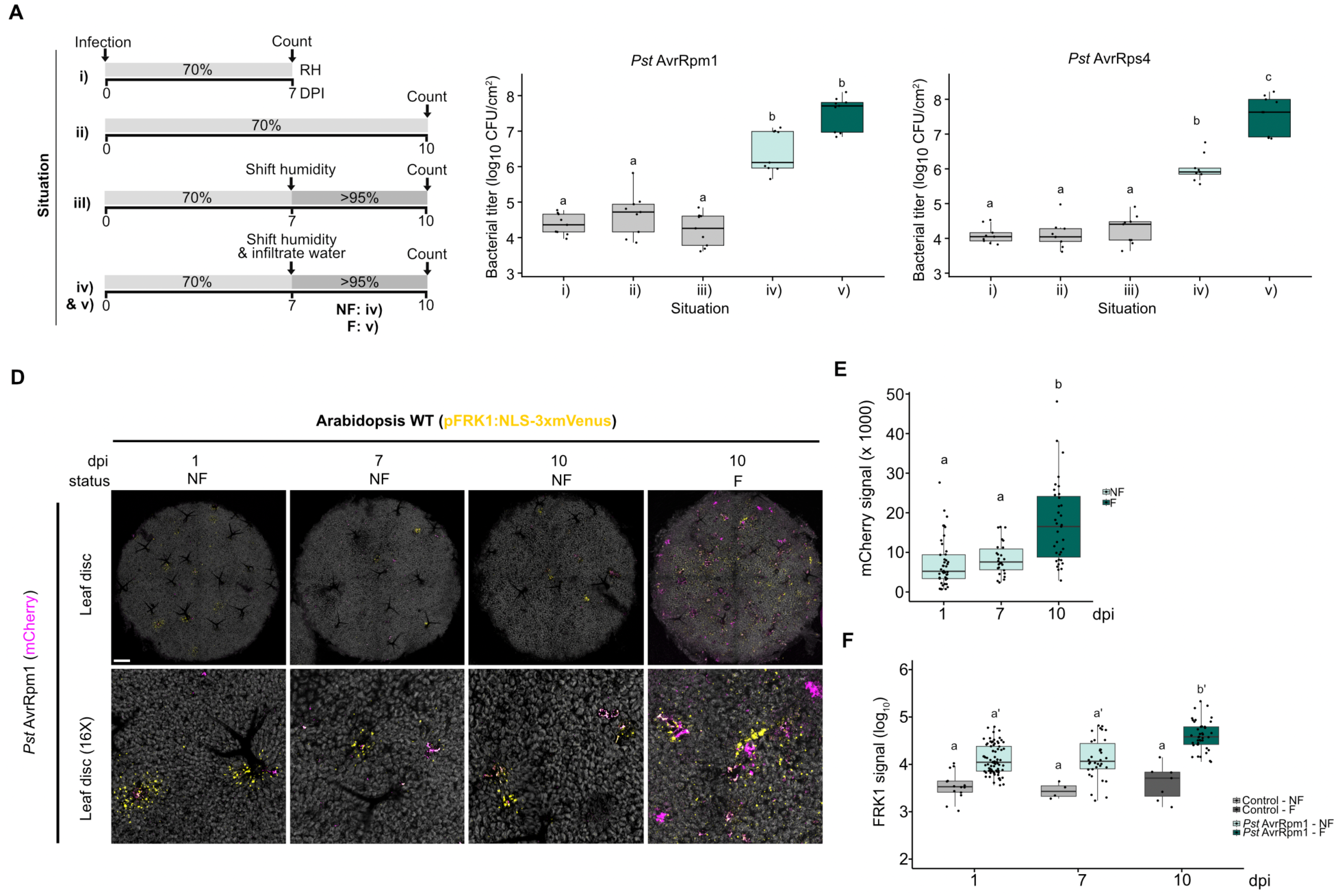
Apoplast hydration re-initiates microbial growth despite ETI pre-activation. (**A**) Schematic of the experimental setup to evaluate a change in bacterial growth in response to humidity and/or apoplastic water in four-week-old Arabidopsis plants. In Situation **I-V**, all plants were inoculated at day 0 with *Pst* AvrRpm1 or *Pst* AvrRps4 (1 x 10^5^ CFU/mL) and maintained under 70% RH for the first 7 days, and up to 10 days (Situation **II**). In Situation **III-V**, plants were domed to increase humidity to >95% between day 7 and 10. In Situation **IV** and **V**, leaves were syringe infiltrated with water and maintained in a NF (IV) or F (V) state for the between day 7 and 10. Bacterial counts were monitored at day 7 for Situation **I** and day 10 for Situation **II-V**. (**B-C**) Bacterial populations undergoing ETI reinitiate growth upon water infiltration in the apoplast. Bacterial counts from Arabidopsis leaves infiltrated with *Pst* AvrRpm1 (**B**) or *Pst* AvrRps4 (**C**) monitored as outlined in (**A**). Different letters indicate statistical significance (p < 0.05) after performing a Wilcoxon rank-sum (Kruskal-Wallis) followed by a Bonferroni correction. (**D**) Localized maintenance of immune signaling at bacterial proliferation sites under ETI. Confocal microscopy images from Arabidopsis leaves containing the immune transcriptional reporter transgene pFRK1:NLS-3xmVenus after surface-inoculation by spraying mCherry-tagged *Pst* WT or *Pst* AvrRpm1. (**E**) Average mCherry signal per confocal images, representing bacterial signal as a log_10_ arbitrary unit (AU). Different letters indicate statistical significance (p < 0.05) after performing a Wilcoxon rank-sum (Mann-Whitney U), two-sided with a Bonferroni correction. (**F**) Average pFRK1:NLS-3xmVenus signal per confocal images, representing immunity signal as a log_10_ AU. Different letters indicate statistical significance (p < 0.05) after performing a Wilcoxon rank-sum (Mann-Whitney U) followed by a Bonferroni correction.

Next, we evaluated the plant immune response and *Pst* proliferation in non-flooded and flooded plants at multiple time points. Co-imaging of the Flg22-induced receptor-like kinase 1 (*FRK1*) Arabidopsis immune transcriptional reporter line (*28*) with mCherry-tagged *Pst* AvrRpm1 revealed a small number of hotspots of *FRK1* reporter signal clustering near *Pst* AvrRpm1 in non-flooded conditions over 10 days after initial infection (Fig. 4D-F). However, flooding the apoplast at 7 dpi and evaluating *FRK1* reporter and *Pst* AvrRpm1 signal at 10 dpi showed that bacterial cells started to proliferate to WT-like levels despite strong immune elicitation in surrounding cells (Fig. 4D-F). This observation supports the hypothesis that ETI does not induce cell death at cellular resolution but simply prevents microbial growth at the infection site (*29-30*). Similar results were obtained when plants were first flooded with water and then sprayed with bacterial inoculums, which represents a more natural infection route for bacteria (Fig. S13). These results suggest that apoplastic water in and of itself is sufficient to prevent the execution of immune functioning against bacteria, without interfering with immune signaling.

### Microbial stasis is caused by a desiccation-induced osmotic stress

If microbial stasis is driven by a lack of apoplastic water availability, we hypothesized that *Pst* WT would experience growth arrest in conditions that are unfavorable to water-soaking induction. We found that *Pst* WT levels were similar to those observed of *Pst* AvrRpm1 or *Pst* AvrRps4 after three days in the apoplast of plants exposed to relative humidity (RH) levels of 40% (Fig. 5A). This is in sharp contrast to the strong growth that occurs at >95% RH, conditions in which water-soaking develops with *Pst* WT (Fig. 5A). In contrast, the growth of avirulent *Pst* strains was unaffected by humidity levels (Fig. 5A). This is consistent with a lack of apoplastic hydration during ETI as shown here (Fig. 5B) and as previously reported (*13*).

**Figure 5:**
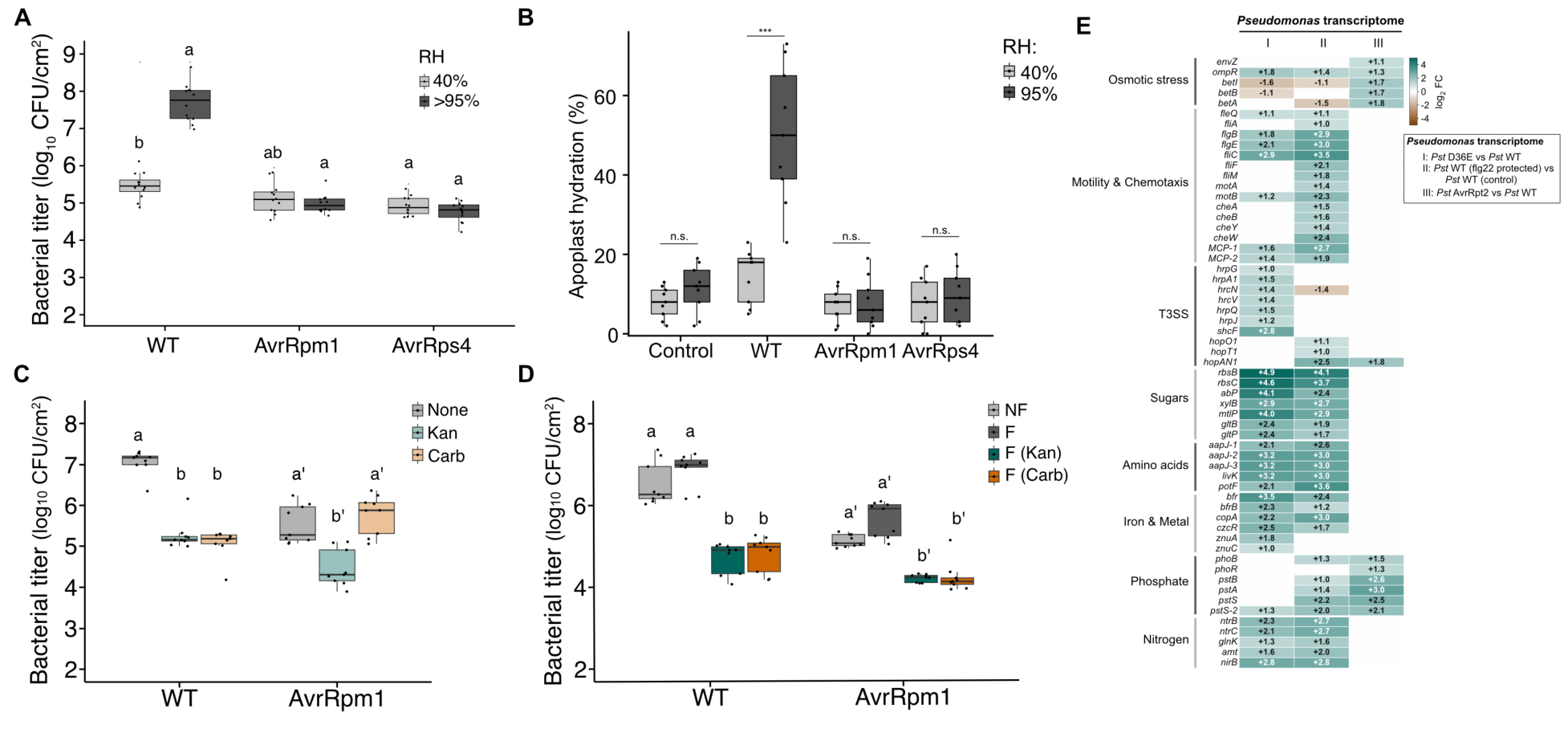
Effector-triggered immunity induces an osmotic stress that drives stasis in a bacterial pathogen. (**A**) Impact of humidity on effector-triggered susceptibility and effector-triggered immunity. Arabidopsis leaves were infiltrated with *Pst* WT, AvrRpm1 or AvrRps4 (1 x 10^5^ CFU/mL). Plants were kept at either 40% or >95% relative humidity (RH) levels. Bacterial titers were monitored at 3 dpi. Different letters indicate statistical significance (p < 0.05) after performing a Pairwise Wilcoxon test followed by a Bonferroni correction. (**B**) ETI prevents hydration of the apoplast during an infection. Apoplastic hydration was measured 24 hpi post infiltration of a control solution (10 mM MgCl_2_), *Pst* WT, AvrRpm1 or AvrRps4 (1 x 10^5^ CFU/mL). Plants were placed at either 40% or >95% RH levels over the course of the experiment. Asterisks indicate statistical significance (p < 0.0005) between RH levels after Wilcoxon rank-sum test (Mann-Whitney U) on individual inoculations. (**C**) Bacterial cell division, but not metabolism, is arrested during ETI. Arabidopsis leaves of pUBQ10-RCI2A-tdTomato, which possess a kanamycin resistance gene, were infiltrated with *Pst* WT or AvrRpm1 (1 x 10^5^ CFU/mL). A control solution (water), kanamycin (50 μg/ml) or carbenicillin (400 μg/mL) was infiltrated, as indicated, into the same leaves 3 days post inoculation. Bacterial population levels were measured 6 hours post infiltration of the antibiotics. Different letters indicate statistical significance (p < 0.05) after performing a Kruskal-Wallis with Dunn’s post-hoc test (Bonferroni) for *Pst* WT and a one-way ANOVA followed by a Tukey’s HSD test for *Pst* AvrRpm1. (**D**) Breaking osmotic stress re-initiates bacterial sensitivity to a cell division-targeting antibiotic. Arabidopsis leaves were infiltrated with *Pst* WT or AvrRpm1 (1 x 10^5^ CFU/mL). Water was infiltrated 2 days post inoculation and leaves were kept in either a NF or F state for a total of 6 hours. The F state leaves were then allowed to revert to the pre-infiltration state. All leaves were then re-infiltrated with water, kanamycin (50 μg/mL) or carbenicillin (400 μg/mL), as indicated and allowed to return to pre-infiltration state. Bacterial populations were measured 6 hours after the final treatment (with or without antibiotics). Asterisks indicate statistical significance (***p < 0.0005) after performing a Kruskal-Wallis followed by a Dunn’s post-hoc (Bonferroni) test. (**E**) ETI specifically-induces an osmotic stress response in *Pst*. Heatmap (log_2_ fold change gene expression) of key selected genes in the indicated *Pst* strain associated with bacterial osmotic stress, virulence, metabolism and nutrient acquisition responses from a previously published RNA-seq dataset (31). Situation I) Comparison of an effector-less *Pst* strain (D36E; 36 effectors deleted) relative to *Pst* WT, II) *Pst* WT in leaves that were previously protected by a flg22 treatment compared to *Pst* WT in unprotected leaves and III) *Pst* AvrRpt2 compared to *Pst* WT.

To test if ETI prevents pathogenesis by triggering a desiccation-induced microbial stasis, we inoculated virulent and avirulent strains of *Pst* into Arabidopsis leaves, followed by treatment with antibiotics that allow for the distinction between bacteria that are actively dividing or not (*27*). As expected, levels of viable *Pst* WT were strongly reduced by treatment with an antibiotic that selectively targets actively dividing cells (carbenicillin), as well as by kanamycin, an antibiotic that targets dividing and non-dividing cells (Fig. 5C). In contrast, bacteria experiencing ETI (*Pst* AvrRpm1) were not affected by carbenicillin, but were affected by kanamycin (Fig. 5C). This selective antibiotic sensitivity suggests that *Pst* AvrRpm1 remains viable but does not actively divide in leaves undergoing ETI. Next, we hypothesized that if ETI-associated osmotic stress causes stasis in bacteria, then re-hydrating the apoplast prior to antibiotic treatment, should increase bacterial sensitivity to carbenicillin reversing the osmotic stress response. In contrast to the situation seen in non-flooded leaves (Fig. 5C), flooding leaves with water for six hours before antibiotic treatment resulted in *Pst* AvrRpm1 and *Pst* AvrRps4 becoming equally susceptible to carbenicillin as to kanamycin (Fig. 5D). This result indicates that bacterial metabolism and division were re-initiated by the addition of water in the apoplast.

We analyzed previously published transcriptomic datasets of *Pst* experiencing different immune responses (PTI or ETI) in Arabidopsis WT and immune mutant plants (*31*) to gain further insight into the impact of ETI as an osmotic stress inducing mechanism. Indeed, we find that *Pst* AvrRpt2 induces the expression of its glycine betaine biosynthetic operon (*betA*, *betB*, *betI*) via a two-component system involved in osmotic stress sensing (EnvZ/OmpR) when compared to *Pst* WT (Fig. 5E). Glycine betaine was previously shown to be a key osmoprotectant produced by *Pseudomonas syringae* in response to osmotic stress *in vitro (32).* Further analysis of *Pst* AvrRpt2 transcriptomes in Arabidopsis mutants for genes involved in the recognition of AvrRpt2, SA biosynthetic or signaling mutants, as well as ethylene and jasmonic acid biosynthetic mutants further revealed a specific role for SA signaling in mediating this osmotic response in *Pst* (Fig. S14). Loss of SA signal transduction (*npr1*) reduced osmotic stress on *Pst* AvrRpt2 to levels nearing those found in the Arabidopsis *rpm1/rps2* mutant, which lacks the NLRs that recognize AvrRpt2 (Fig. S14). Interestingly, osmotic-stress responsive genes up-regulated in *Pst* AvrRpt2 were either down-regulated or unaffected in *Pst* strains that were under strong plant PTI responses (Fig. 5E). Instead, an effector-less *Pst* strain and a WT *Pst* inoculated in plants that were previously protected by a flg22 treatment displayed strong expression of genes associated with nutrient transporter and chemotaxis/motility (Fig. 5E). Alongside a previously reported reduction in global activity (ribosomal and housekeeping genes being negatively affected by PTI) (*31, 33*), this suggests that PTI functions rather as an extracellular starvation-inducing mechanism. Together, these results highlight a previously underappreciated dichotomy between PTI and ETI on bacteria, where a strong PTI response triggers a nutritional stress which activates a scavenging behavior in *Pst* while ETI induces a strong osmotic stress. While diverging mechanistically, both PTI and ETI appear to control microbial proliferation by reducing the quality of the extracellular niche environment.

## Discussion

A large body of work exists pertaining to the unravelling of how immune receptors are activated, assembled and signal inside plant cells (*34, 35*), which in turn has guided disease resistance engineering efforts (*36-40*). However, our current understanding falls short of explaining why immune activation leads to microbial growth arrest. At least five non-mutually exclusive hypotheses for how immunity achieves this can be envisaged: that immunity triggers cell death to restrict microbial access to host tissue; that it spatially confines pathogen spread; that it reduces the extracellular environment quality; reduces virulence by impacting effector delivery or that it directly affects microbial viability (*13, 22, 30, 41-44*). Our findings bear on each of these hypotheses and converge on a unifying model in which PTI and ETI impose a hostile apoplastic environment, through desiccation and nutrient limitation, sufficient to arrest bacterial growth independently of cell death. Plant immunity has long been understood through the lens of protein biochemistry. The implication that extracellular water alone is sufficient to inhibit effective execution of plant immunity without altering biochemical functions or immune signaling exemplifies the need to link immunity-driven biochemical events to physiological outcomes.

The relationship between ETI, the hypersensitive response, and disease resistance has long been an area of interest. Resistance following ETI induction spans a spectrum from non-HR to HR-associated reactions depending on multiple factors relating to pathogen type, inoculum and the nature of the recognition by the plant (*26*). HR-associated responses likely represent the extreme end of an ETI intensity spectrum rather than a mechanistically distinct outcome. Consistent with the HR phenotype intensity, we show that the degree of bacterial growth arrest directly correlates with the extent of plant tissue desiccation during ETI (Fig. 3B). Bacteria experiencing strong PTI appear to actively upregulate nutrient acquisition systems, a response that is similar to, but weaker than, that observed during ETI. This raises the possibility that immune intensity exists on a continuum ranging from mild nutritional stress at the PTI end to overt osmotic stress-driven growth arrest at full ETI, providing a parsimonious explanation for the graded nature of immunity-mediated bacterial growth inhibition. High-dose infiltration of avirulent strains has been shown to suppress SA signaling at the epicenter of the infection zone while leaving it intact at the margins (*30*). This pattern is consistent with our observation that high inoculum levels compromise ETI efficacy, allowing for bacterial growth. We speculate that resistance at infection margins reflects reduced effector delivery and diminished host immune suppression as bacterial density declines, rather than a spatially distinct immune mechanism. Critically, we show that localized immunity confines bacterial populations to narrow apoplastic microenvironments (Fig. 4D), yet this restriction is fully reversed by apoplastic water supplementation. This demonstrates that growth inhibition, rather than spatial confinement, is the primary mechanism by which ETI restricts bacterial proliferation. We speculate that the local confinement previously reported may be caused by bacterial osmotic stress preventing growth locally. Furthermore, our results suggest that different types of disease resistance induce degrees of water immunity. That is, PTI results in a physiological state where bacteria cannot induce water soaking (*4, 45, 46*). Non-HR inducing ETI appears to induce a similar state, whereas HR-inducing ETI appears to result in a state wherein stomata remain open, but additional mechanisms must also exist that result in a rapid loss of water from infected tissues (Fig. 3 and Fig. S10)

Bacterial stasis has been observed in long-term experiments in Arabidopsis plants inoculated with non-pathogenic members of the leaf microbiome, non-virulent and avirulent *Pseudomonas* strains (*27*). Whether this stasis could be reverted despite strong immunity activation was unknown. Nutrient and water depletion during immune activation has previously been reported (*13, 42, 47*). The extent to which these stresses were the causal agents behind disease resistance and stasis was unclear, despite *in vitro* evidence that osmotic stress experienced by avirulent *Pst* strains could be growth inhibiting (*48*). Combinatorial antibiotics treatment targeting either bacterial metabolism or active cell division revealed that, during ETI, bacteria still have at least moderate metabolic activity but are not dividing. This was likely driven by the osmotic stress experienced by local cell desiccation, as supported by bacterial transcriptome analysis during ETI.

Our findings establish that plant immunity actively remodels the physiological state of invading bacteria, through osmotic stress and nutrient restriction, as a mechanism of disease resistance, distinct from and complementary to intracellular immune signaling. Given that the apoplast is a nutrient-poor environment even in the absence of infection, and that niche remodeling by the host is likely a general feature of plant-pathogen interactions, it will be important to determine whether host-imposed apoplastic control of pathogen physiology represents a conserved resistance mechanism across diverse plant-pathogen systems. More broadly, our results argue that a complete understanding of plant immunity requires examining its consequences from the pathogen’s physiological perspective, not only from the host’s intracellular signaling networks. Extending this framework to crop pathosystems may reveal new physiological vulnerabilities in microbial pathogens that could be exploited for disease control.

## Author contributions

Conceptualization: CRL, PM

Methodology: CRL, SG, PM

Investigation: CRL, SG, YH, SM, FGL, JZ, AP, AR

Visualization: CRL, SG, YH, JZ

Funding acquisition: PM, XFX, GC, LN

Project administration: CRL, PM

Supervision: CRL, XFX, GC, LN, PM

Writing – original draft: CRL, SG, PM

Writing – review & editing: CRL, SG, YH, JZ, SM, XFX, LN, GC, PM.

## Acknowledgements

The authors would like to thank Sheng Yang He and He lab members for the insightful discussions regarding the findings in this study. This study was supported by a Natural Sciences and Engineering Research Council of Canada (NSERC) Discovery Grants to P.M. C.R.-L. was supported by a Quebec Doctoral Graduate Scholarship (FRQ-NT), EMBO postdoctoral fellowship and Swiss National Science Foundation postdoctoral fellowship. A.P. was supported by an NSERC Doctoral Graduate Scholarship. This work was in part supported by a PhD grant from the French ministry of higher education and research to S.M, a grant from the Agence Nationale de la Recherche NEPHRON project (ANR-18-CE20-0020-01) to L.D.N. and was set within the framework of the ‘Laboratoires d’Excellences’ (LABEX) TULIP (ANR-10-LABX-41) and of the ‘Ecole Universitaire de Recherche’ (EUR) TULIP-GS (ANR-18-EURE-0019). G.C. and J.Z. were supported by John and Joan Fiddyment Endowed Chair funds to G.C.

## Supplementary Materials

Materials and Methods

Figs. S1 to S14

References (1–48)

## Material and methods

### Plant material and growth conditions

For most experiments conducted in this study (U. de Sherbrooke), Arabidopsis plants were grown in PromixTM soil (PremierTech) in growth chambers with a 12 h light/dark photoperiod, with relative humidity of ∼60% at 21 °C. Light intensity was measured at 180 µmoles/m^2^/s when lights were on. Four– to five-week-old Arabidopsis plants were used for all experiments described herein, unless stated otherwise.

For experiments associated with Figure S7, cauliflower plants (*Brassica oleracea* cv. *botrytis* var. *Clovis* F1, Vilmorin) were grown under glasshouse conditions for five weeks. After inoculation, all plants were placed in a growth chamber at 22°C in short-day conditions (8h light, 190 µmol/m²/s) and 70% relative humidity.

For confocal imaging experiments (Fig. 4D-F and Fig. S13), Arabidopsis plants were grown in Sunshine Mix #1 soil (Sun Gro Horticulture) in growth chambers with a 10-h light/14-h dark photoperiod (100 μM m^−2^ s^−1^), with relative humidity of ∼70% at 23 °C.

### Bacterial disease assays

Pseudomonas syringae pv. tomato DC3000 WT and mutant strains were cultured overnight at 28 °C in Luria-Bertani (LB) media containing 50 mg/L of rifampicin. On the day of the infection, fresh LB media was inoculated with 0.5 mL of the overnight culture and bacteria were collected when OD_600_ reached between 0.8–1. Bacteria were centrifuged for 10 min at 4000 × g and the pellet resuspended in ddH_2_O. Bacterial density was adjusted to 0.2 (1 × 10^8^ CFU/mL) prior to further dilutions. Bacterial infections were carried out between 14:00–15:00 (zeitgeber time of 6:00–7:00).

All inoculations of Arabidopsis leaves were performed by syringe-infiltration. Infiltrated plants were all kept under ambient humidity levels for 1 h to allow water to evaporate (or not in the F state), then domed with a plastic unit to maintain high humidity (>95% RH).

Bacterial growth in planta was monitored by harvesting infected Arabidopsis leaves, surface sterilizing in 80% ethanol and rinsing in sterile water twice. Leaf disks were taken from three leaves from the same plant (one per leaf; total of three leaf disks) using a cork borer (6 mm in diameter) and ground in sterile 10 mM MgCl_2_. Three biological replicates were performed for each experiment. Colony-forming units (CFU) were determined by making serial dilutions (10^0^-10^-6^) and plating on LB plates containing 50 mg/L of rifampicin. Each dilution was plated in three technical replicates.

*Xanthomonas campestris* pv. *campestris* CN06 WT (*49*) and mutant CN06*ΔxopJ6 (20)* were cultivated for 60 hours on solid MOKA media containing 50 mg/L of rifampicin. On the day of infection, bacteria were centrifuged for 10 minutes at 6000 x g and washed twice in 10 mM MgCl_2_. Bacterial density was adjusted to 0.0001 (5 × 10^4^ CFU/mL). Bacterial infections were carried out between 8:30–9:30 am. All inoculations of cauliflower plants were performed by syringe infiltration on the third leaf. Infiltrated plants were all kept under ambient humidity levels for 1.5 day to allow HR development. On the day following inoculation, some plants were sealed with silicon grease between 16:30–17:30 (at dusk), and all plants were then domed with a plastic lid to maintain high humidity (>95% RH). Bacterial growth *in planta* was monitored by harvesting cauliflower leaf disks using a cork borer (6 mm in diameter). Leaf disks were taken from the same leaf (one per strain) and two or three biological replicates were performed for each experiment. After grinding in 10 mM MgCl_2_, colony-forming units (CFU) were determined by making serial dilutions (10^0^-10^-6^) and plating on MOKA plates containing 50 mg/L of rifampicin and 30 mg/L of pimaricin. Each dilution was plated in three technical replicates.

### Callose deposition imaging and quantification

Leaves of four-week-old Arabidopsis plants were infiltrated as described in the associated figures. Leaf disks were taken from four leaves from the same plant (one per leaf; total of four-leaf disks) using a cork borer (6 mm in diameter) and placed in 24-well plates containing 95% ethanol. Plates were placed on a rotating shaker and ethanol renewed three times over the course of four hours to allow leaf decolouration. Once leaf disks were decoloured (white), they were rinsed in 67 mM K_2_HPO_4_ for 1 hour. K_2_HPO_4_ was removed and replaced by callose staining solution (K_2_HPO_4_ 67mM containing 0.1% aniline blue) for another hour. The staining solution was then removed, and leaf-disks quickly rinsed in 67 mM K_2_HPO_4_. The rinsing solution was then removed and replaced with fresh 67 mM K_2_HPO_4_ solution and placed back on the rotating shaker. The plate containing leaf disks was then placed in a 4°C refrigerator until imaging.

Callose deposition was imaged using a FV-3000 Olympus confocal microscope. Callose quantification was measured using a reversed-color analysis function of ImageJ software and quantifying the amount of callose deposition in terms of pixels against the total amount of pixels found in each image, which resulted in a percentage of callose area per image.

### Trypan blue staining

Leaves of four-week-old Arabidopsis infected with Pst AvrRpt2 were harvested at 12h or 24h post inoculation and boiled in staining solution (10 ml lactic acid, 10mL glycerol, 10mL water-saturated phenol, 0.1g trypan blue, dissolved in 10mL ddH2O) for 2min. Then the leaves were incubated in the staining solution for another 1h with shaking. The leaves were de-stained in 2.5 g/ml chloral hydrate solution for several times until the background was completely colorless and transferred to 70% glycerol for subsequent imaging.

### Quantification of apoplastic reactive oxygen species

Leaf disks from four-week-old Arabidopsis plants were collected using a 4 mm diameter biopsy punch and placed into white 96-well plates (Corning) containing 100 μL of distilled water for 16 h (overnight). Prior to ROS quantification, water was removed and replaced with ROS assay solution (100 μM Luminol [Millipore-Sigma], 20 μg/mL horseradish peroxidase [Millipore-Sigma]), with or without immune elicitors. Light emission was measured using a TECAN Spark® plate reader. Experiments were repeated three times.

### Peroxidase activity assay

Single leaves were excised from Arabidopsis WT plants. Leaf disks (diameter 5 mm) were taken from each leaf and were washed for 1 h in 1 mL of ½ MS solution, with agitation. After washing, disks were carefully transferred to individual wells of a clear 96-well assay plate, using forceps while minimizing damage. Each well received 50 μL of ½ MS solution with vehicle control (H_2_O) or was supplemented with flg22 peptide. Plates were sealed with Parafilm and were incubated for 20 h at room temperature with agitation. Leaf disks were removed and each well received 50 μL of a 1 mg/ml solution of 5-aminosalicylic acid, pH 6.0, with 0.01% hydrogen peroxide, using a multichannel pipette to minimize timing differences. The reaction was allowed to proceed for 3 minutes and was stopped by the addition of 20 μL of 2 N NaOH, using a multichannel pipette in the same order as above, to ensure equal time for enzymatic reaction in each well, prior to reading the OD_600_ on an TECAN Spark® plate reader.

### MAPK phosphorylation assay

Arabidopsis leaves were infiltrated with water and kept in a NF or F state prior to collecting leaf punches (5 mm) and carefully transferring individual punches to a clear 96-well assay plate. Leaf disks were kept in sterile ddH_2_O for 16 hours to allow tissues to recover from the punching stress. Leaf disks were spiked with a flg22 solution at a final spiked concentration of 1 μM. Plant material was collected and flash-freeze in liquid nitrogen at 0, 15 or 30 minutes post elicitation. Tissues were grounded in protein extraction buffer (0.35 M Tris-HCl pH 6.8; 30% (v/v) glycerol; 10% (v/v) SDS; 0.6 M DTT; 0.012% (w/v) bromophenol blue) and loaded unto SDS-PAGE gel for Western-blotting. An anti-MAPK p44/42 antibody from Cell Signaling Technology (#9101S) was used to evaluate MAPK phosphorylation status. Coomassie blue staining was done on the membrane post anti-MAPK p44/42 evaluation to assess that protein loading was similar across samples.

### Apoplast antibiotic treatments

For antibiotic impact on virulent or avirulent *Pst* strains *in planta*, *Pst* WT was transformed with an empty vector (pOT1e) or a vector carrying the avirulent gene AvrRpm1 (pOT1e-AvrRpm1). Both vectors carried tetracycline resistance cassettes as not to interfere with other antibiotic resistance profiles. Kanamycin (50 μg/mL) or carbenicillin (400 μg/mL) was infiltrated in Arabidopsis leaves carrying a kanamycin resistance cassette to reduce the impact of kanamycin on plant tissues undergoing an infection with either *Pst* WT or *Pst* AvrRpm1. Experiments were carried as further described in Figure 5C or 5D.

### Quantification of leaf water content by hyperspectral imaging

For plant phenotyping and water content parameters analysis, plants were transferred from growth chambers to a PlantScreen™ phenotyping system (Photon Systems Instruments, Drásov, Czechia) at the Eastern Canadian Plant Phenotyping Platform (Sherbrooke). First, RGB images were acquired using a plant screen to isolate plant-associated pixels from background pixels. Second, SWIR (short-waved infrared) images were acquired to perform water content analysis.

Pictures were analyzed using the phenotyping system program’s BilReader. Plant masks were manually assigned to each area to be quantified either the whole plant for pictures and spray inoculations or separated leaves for infections. Water content was quantified using the WATER1 formula R1440/R990 provided by Photon Systems Instrument, which is essentially a ratio of water reflectance values. Plots were generated using the mean area and standard deviation and representative false-color images based on the mean area were generated using the same scale for all plants within the same experiment.

### Confocal microscopy imaging

For Figure 4D, mCherry labelled *Pst* DC3000(AvrRpm1) (Tn*7*-3xmCherry) was infiltrated (OD_600_=0.0002) on 4-week-old *FRK1* transgenic plants (*pFRK1-NLS-3xmVenus*) growing in soil. Three plants were inoculated for each treatment at each time point. Infiltrated plants were wiping extra water at the bottom side away and kept under ambient humidity levels for 1-2 h to allow water to evaporate and let plant leaves return to a pre-infiltration appearance. Then plants were domed with a plastic lid to maintain high humidity (>95% RH) for 4 h. Then image at 1 dpi and 7 dpi. At 7 dpi, the inoculated plant leaves were infiltrated with water and domed with a plastic lid to maintain high humidity (>95% RH) for 3 days and 1 hour. Image at 10 dpi. Bacterial population was measured at 0, 1, 7, and 10 dpi. Confocal imaging was performed on a Leica TCS SP8 microscope. Pictures were taken with a 5x immersion objective for tile-scan with 10% overlap. The following excitation and emission parameters were used for different fluorophores: mVenus488 nm, 493 – 540 nm; mCherry 552 nm, 586 – 635 nm; chlorophyll 638 nm, 650-720 nm. Sequential scanning was applied to avoid fluorescence interference between channels. All images were taken under identical settings (lens, laser power, pinhole size, detector gain and interval of Z stack) for comparison of fluorescence intensity over time. ImageJ was used to quantify fluorescence intensity at each infection site for all images. A fixed threshold was applied consistently to each fluorescence signal. At least six images from three different plants were analyzed for each treatment.

### RNA extraction and sequencing

Arabidopsis leaves were harvested and immediately flash-freezed in liquid nitrogen for further RNA extraction protocol. RNA was extracted from flash-frozen, ground leaf tissue followed with QIAZOL (QIAGEN) reagents followed by on-column DNase treatment (QIAGEN), according to the manufacturer’s protocol. RNA integrity was evaluated by an Agilent Bioanalyzer 2100 with the Eukaryote Total RNA Nano Series II. cDNA libraries were generated using NEBNext Ultra II Directional RNA Library Prep Kit for Illumina kit (New England Biolabs, USA) according to the manufacturer’s protocol. cDNA libraries were sequenced by RNA-seq at the Université de Sherbrooke RNomics Platform using an Illumina NextSeq 500 system. Approximately 20 million reads were generated per sample.

Reads quality was assessed using FastQC and low-quality sequences removed by using cutadapt with a quality cutoff of 30. The resulting reads were mapped onto the Arabidopsis thaliana genome (TAIR10) using RNA STAR. Mapped reads were counted using featureCounts. Differential gene expression analysis was performed by using the DESeq2 package. A cutoff of q-value <0.01 and absolute log_2_ fold change > 1 was applied to identity DEGs. RNA-sequencing raw data are available at EMBL-EBI ArrayExpress under the identifier E-MTAB-17058.

### Apoplast hydration measurement

Three fully expanded leaves per plant from three plants were excised and apoplast extracted as previously described, with some modifications (*50*). Briefly, the initial weight of freshly excised leaves was measured before being infiltrated with distilled water and weighed again once leaves were fully saturated with water. Leaves were centrifuged at 4000 RCF in 2 mL microcentrifugation tubes containing glass beads at the bottom to prevent tissue collapse during the centrifugation. Leaves were weighed post centrifugation and apoplast hydration determined as previously described. Experiments were repeated at least three times.

### Statistical analysis

All statistical analyses were performed using the bioinformatic software R. Statistical significance was set at p<0.05. Statistical tests used are stated in figure legends. All statistical test assumptions, such as normality and homoskedasticity, were tested. When not respected, non-parametric equivalent tests were performed. In multiple comparison tests, Bonferroni corrections were used.

