## Supplementary Figures for "Extracellular water withdrawal drives disease resistance in the phyllosphere"

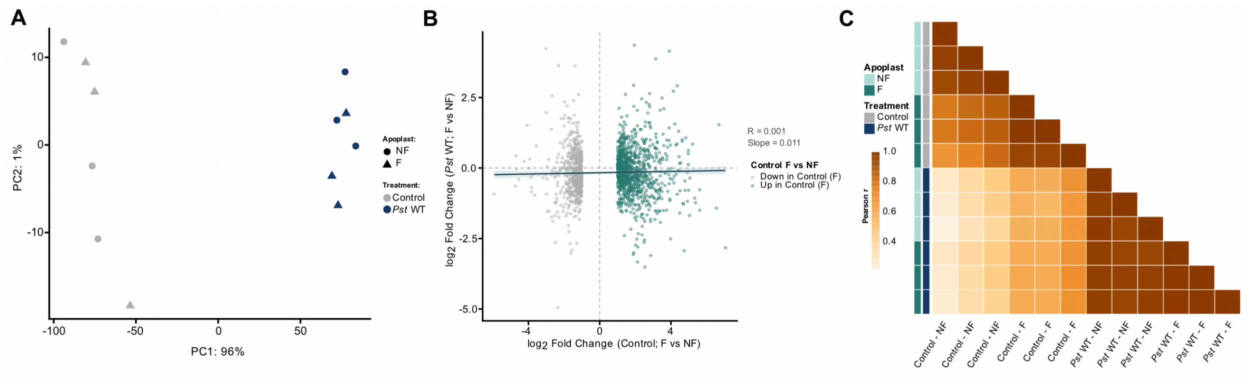

**Figure S1: Transcriptional landscape in Arabidopsis in response to artificial flooding compared to natural water soaking.**

(**A**) Artificial flooding induces minimal differences in the Arabidopsis transcriptome compared to natural water-soaking induced by *Pst* WT. Principal component analysis of transcriptomes from four-week-old Arabidopsis WT plants infiltrated with a control solution (10 mM $MgCl_2$) or *Pst* WT (1 x $10^7$ CFU/mL) at 24 hours post infiltration. Leaves were kept under flooded (F) and non-flooded (NF) apoplast conditions. PCA was performed on variance-stabilized (VST) expression values of the 500 most variable genes across 12 samples (3 biological replicates per condition). Each point represents one biological replicate. (**B**) Flood-induced $\log_2$ fold changes are suppressed in *Pst* WT-infected plants. Scatter plot comparing $\log_2$ fold changes of the 1,714 Control flood-responsive genes (padj < 0.05, $|\log_2 FC| > 1$ in Control F vs NF) in the Control F vs NF contrast (x-axis) against their $\log_2$ fold changes in the *Pst* WT F vs NF contrast (y-axis). Points are coloured by their direction of regulation in Control (teal: upregulated; grey: downregulated). The black line represents the ordinary least-squares linear regression fitted across all genes. $R^2 = 0.001$ and slope = 0.011 indicate that genes strongly regulated by flooding in a pathogen-free context show no coherent directional response to flooding in *Pst* WT-infected tissue, consistent with effector-mediated suppression of the flood-induced transcriptional program. (**C**) Pairwise Pearson correlation coefficients computed on VST-normalized expression values of the 1,714 genes differentially expressed between flooded (F) and non-flooded (NF) apoplast conditions in Control plants (padj < 0.05, $|\log_2 FC| > 1$). Samples are annotated by treatment (grey: Control; navy: *Pst* WT) and apoplast condition (light teal: NF; dark teal: F). Hierarchical clustering (complete linkage) was performed on the correlation matrix. Correlation values are displayed within each cell.

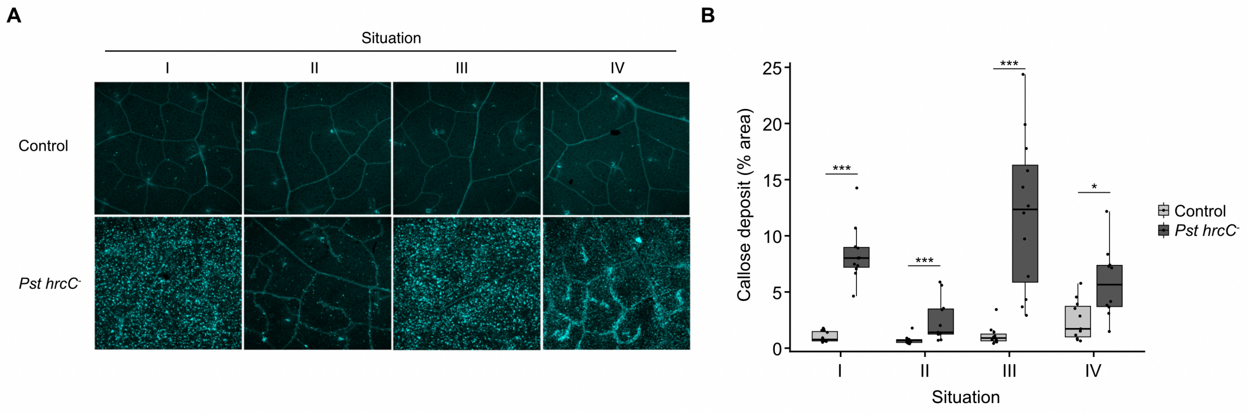


**Figure S2: Shifting the apoplastic state from flooded to non-flooded re-initiates callose deposition by PTI.**

(**A**) Callose deposition inhibition in the flooded apoplast is reversible upon a change of apoplast hydration status. Callose deposition images of four-week-old Arabidopsis at 24 hours post-infiltration of 10 mM $MgCl_2$ (Control) or *Pst hrcC*$^-$ (1 x $10^8$ CFU/mL). Leaves were subjected to four different conditions throughout the experiment. In panels **I** and **II**, leaves were maintained under >95% RH levels in a NF (**I**) or F (**II**) state throughout the entire course of the experiment. In panels **III** and **IV**, leaves were maintained in either a NF (**III**) or F (**IV**) state for the first 16 hours post-infiltration before being shifted to a 70% RH level condition for the remaining 8 hours to allow for apoplastic water evaporation. (**B**) ImageJ quantification of the callose deposition area (% of total area) in the images acquired in (**A**). Asterisks indicate statistical significance (* = p<0.05, *** = p<0.0005) after performing a Wilcoxon rank-sum test (Mann-Whitney U), one test per Situation comparing Control vs *Pst hrcC⁻* independently.

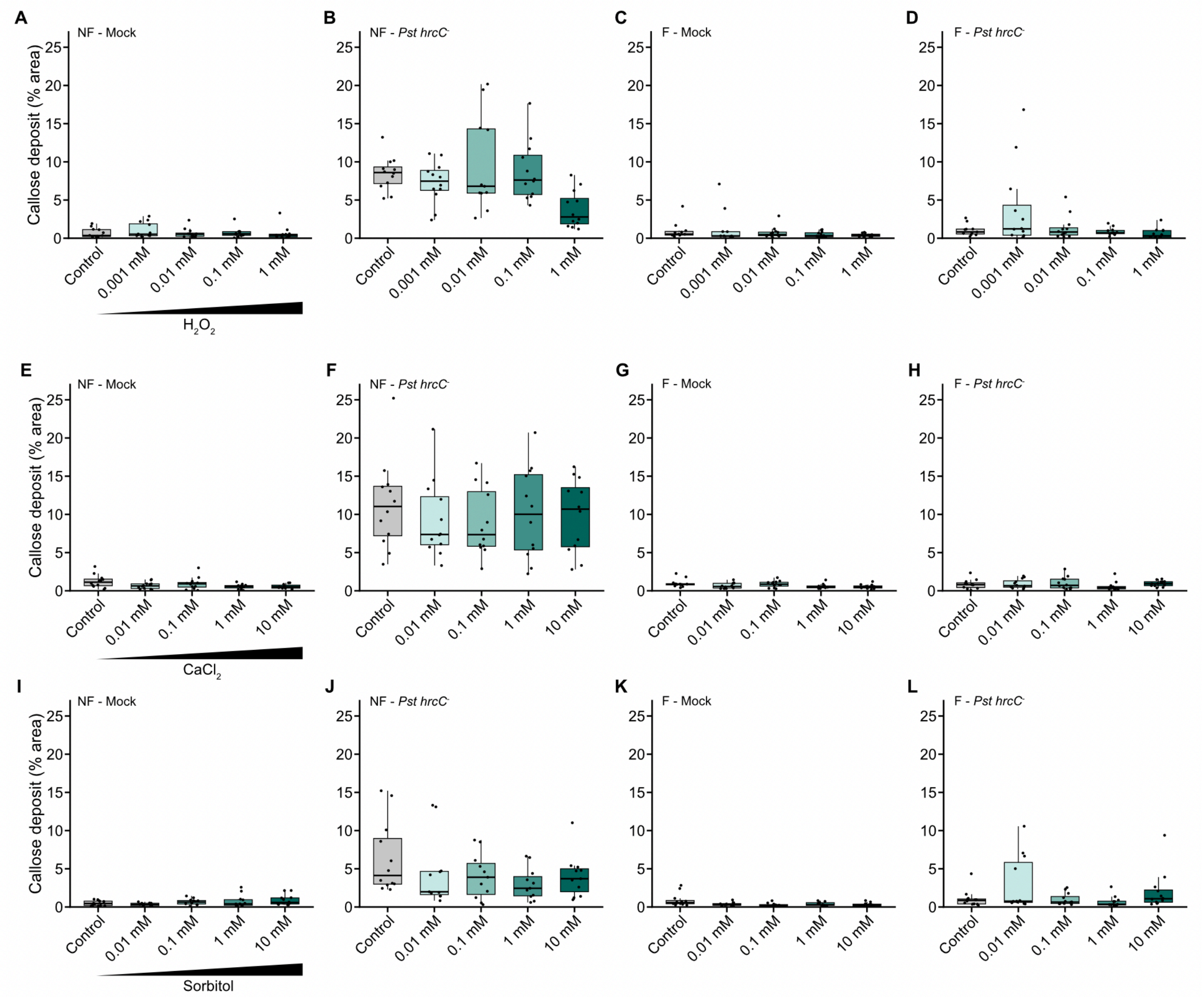


**Figure S3: Callose deposition inhibition under F state is independent of the dilution of ROS species, calcium or reduced osmolarity.**

(**A-D**) Callose deposition quantification in four-wee-old Arabidopsis leaves 24 hours post-infiltration of a water solution (control) with increasing concentrations of $H_2O_2$ (mock), or *Pst hrcC*$^-$ (1 x $10^8$ CFU/L) solution with similar increasing concentrations of $H_2O_2$. The apoplast was either maintained in a NF or F state with these solutions over the course of the experiment. (**E-H**) Callose deposition quantification in four-week-old Arabidopsis leaves 24 hours post-infiltration of a water solution (control) with increasing concentration of $CaCl_2$ (mock), or *Pst hrcC*$^-$ (1 x $10^8$ CFU/mL) solution with similar increasing concentrations of $CaCl_2$. The apoplast was either maintained in a NF or F state with these solutions over the course of the experiment. (**I-L**) Callose deposition quantification in four-week-old Arabidopsis leaves 24 hours post-infiltration of a water solution (control) with increasing concentration of sorbitol (mock), or *Pst hrcC*$^-$ (1 x $10^8$ CFU/mL) solution with similar increasing concentration of sorbitol. The apoplast was either maintained in a NF or F state with these solutions over the course of the experiment.

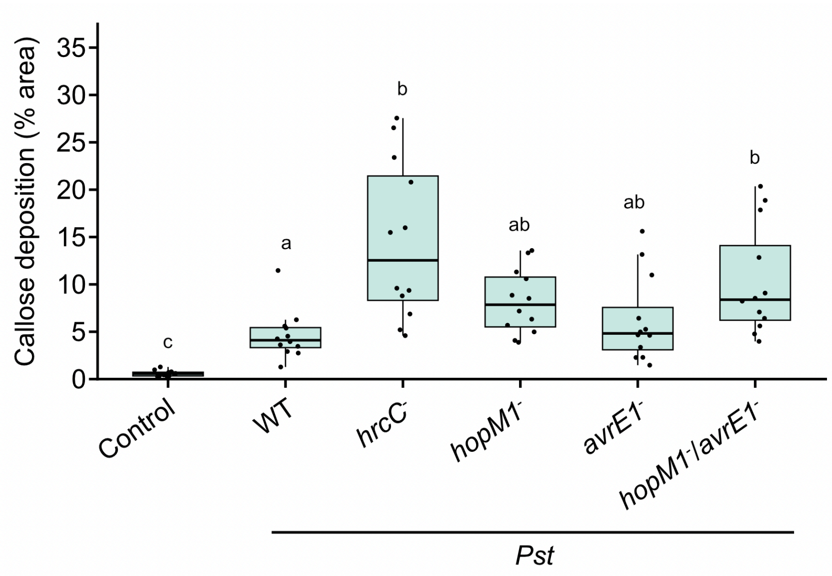


**Figure S4: Callose deposition in Arabidopsis is inhibited by *Pst* water-soaking effectors HopM1 and AvrE1.**

Callose deposition quantification in four-week-old Arabidopsis leaves 24 hours post-infiltration of 10 mM $MgCl_2$ solution (control), WT *Pst*, or mutant strains thereof (1 x $10^8$ CFU/mL), as indicated. Different letters indicate statistical significance ($p < 0.05$) after performing a Kruskal-Wallis followed by a Wilcoxon's post-hoc test (Bonferroni correction).

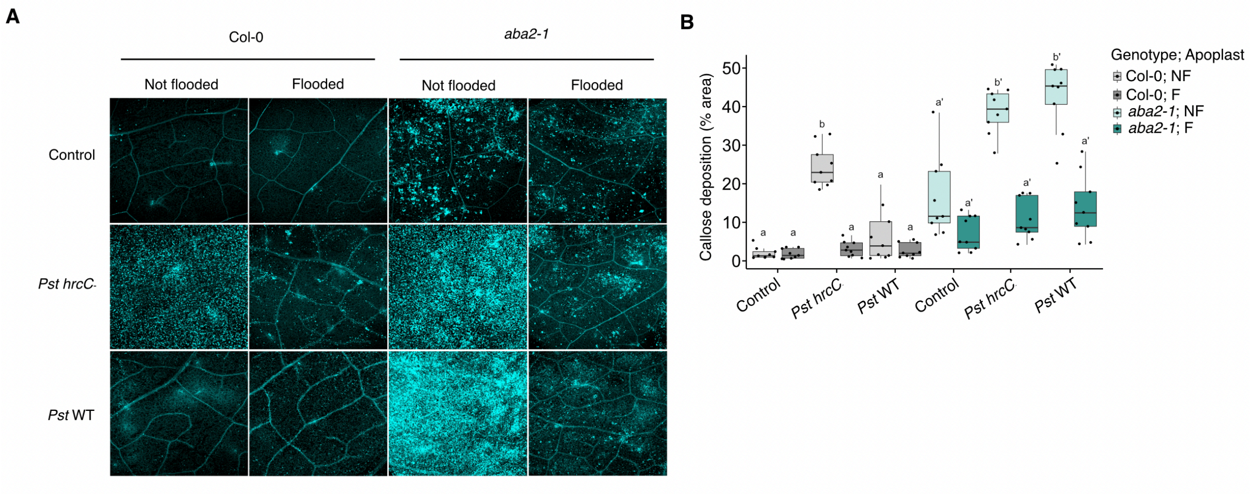


**Figure S5: *Pst*-mediated ABA manipulation is essential to prevent callose deposition via water-soaking induction**

(**A**) Leaves of four-week-old Arabidopsis, WT or *aba2-1*, were infiltrated with 10 mM $MgCl_2$ alone (Control), or with either *Pst hrcC*$^{-}$ or *Pst* WT (1 x $10^8$ CFU/mL), as indicated. Leaves were maintained in a NF or F states and visualized by confocal microscopy 24 hours later. (**B**) ImageJ quantification of the callose deposition area (% of total area) in the images acquired in (**A**). Different letters indicate statistical significance ($p<0.05$) after Pairwise Wilcoxon rank-sum test followed by a Bonferroni correction.

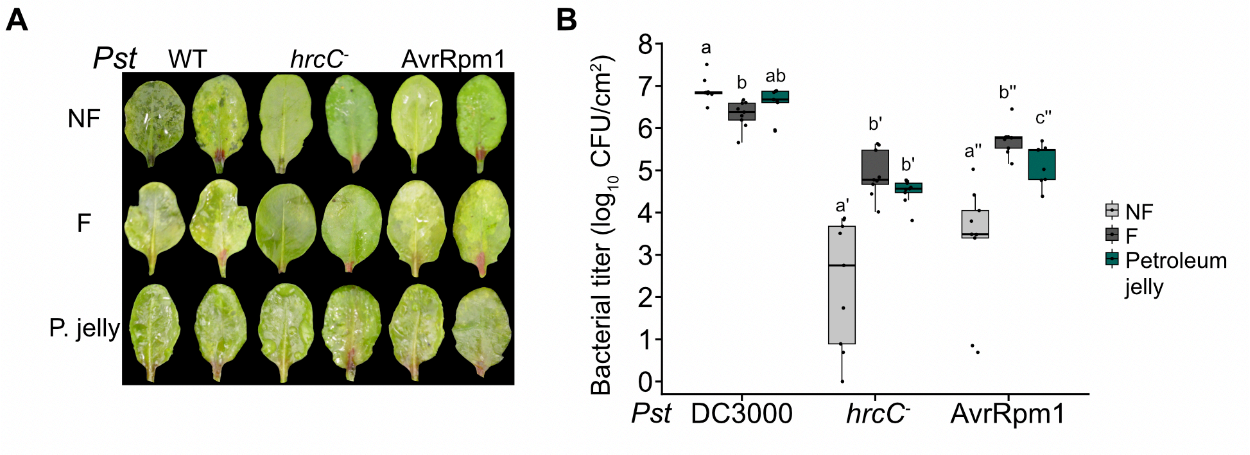


**Figure S6: Leaf sealing with petroleum jelly induces water-soaking-like lesions and drives microbial pathogenesis.**

(**A**) Sealing Arabidopsis leaves with petroleum jelly leads to similar symptoms as those observed by artificial leaf flooding. Photos of four-week-old Arabidopsis WT plants at **24** hours post infiltration with *Pst* WT, *hrcC*- or AvrRpm1 (1 x $10^5$ CFU/mL). Leaves were either kept in a NF state, or F state by syringe infiltrating water (F) or sealing the leaf abaxial side with petroleum jelly (P. jelly). (**B**) Bacterial count at 3 dpi of Arabidopsis leaves infiltrated with the *Pst* WT, *hrcC*- or AvrRpm1 in leaves that were kept in the states mentioned in (**A**). Different letters indicate statistical significance ($p < 0.05$) after performing a pairwise Wilcoxon rank-sum test (Mann-Whitney U) followed by a Bonferroni correction.

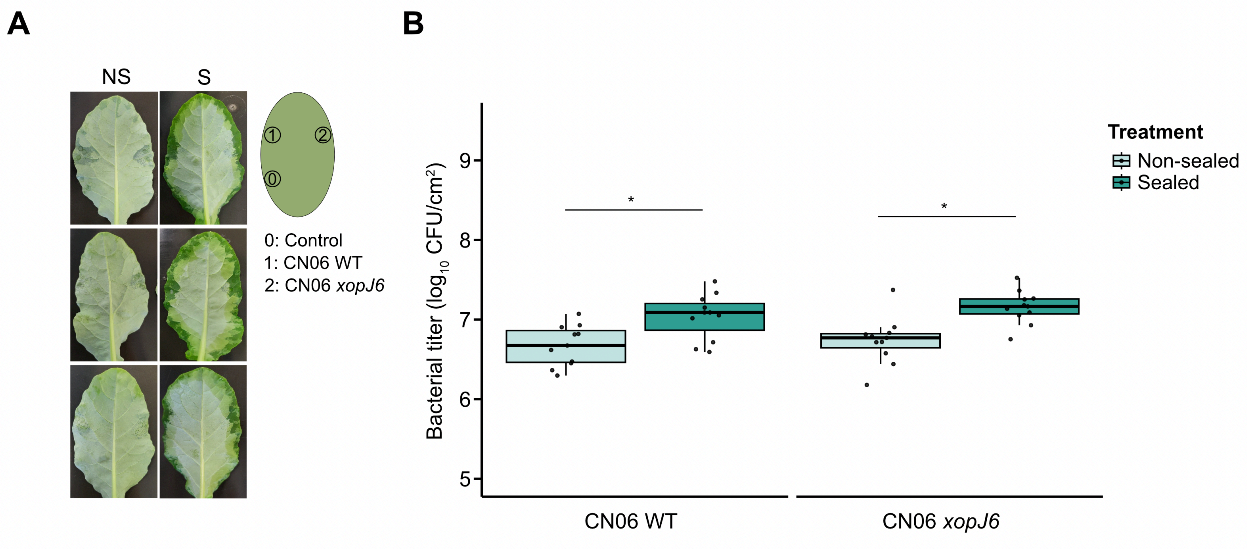


**Figure S7: Impact of leaf margin sealing on virulence of avirulent and virulent *Xanthomonas* CN06 strain.**

(A) Phenotype of cabbage leaves infiltrated with a control solution (10 mM $MgCl_2$), *Xanthomonas* CN06 WT or *xopJ6* mutant (1 x $10^5$ CFU/ml) in which the margin was either sealed (S) or not sealed (NS) with silicone grease at 1.5 dpi. Photos were taken at 3 dpi. (B) Bacterial population in leaves infiltrated with *Xanthomonas* CN06 WT or *xopJ6* mutant (at 5 x $10^4$ CFU/ml), as indicated, at 3 dpi. Asterisks indicate statistical significance ($p<0.05$) after performing a Mann-Whitney U test followed by a Wilcoxon rank-sum post-hoc test.

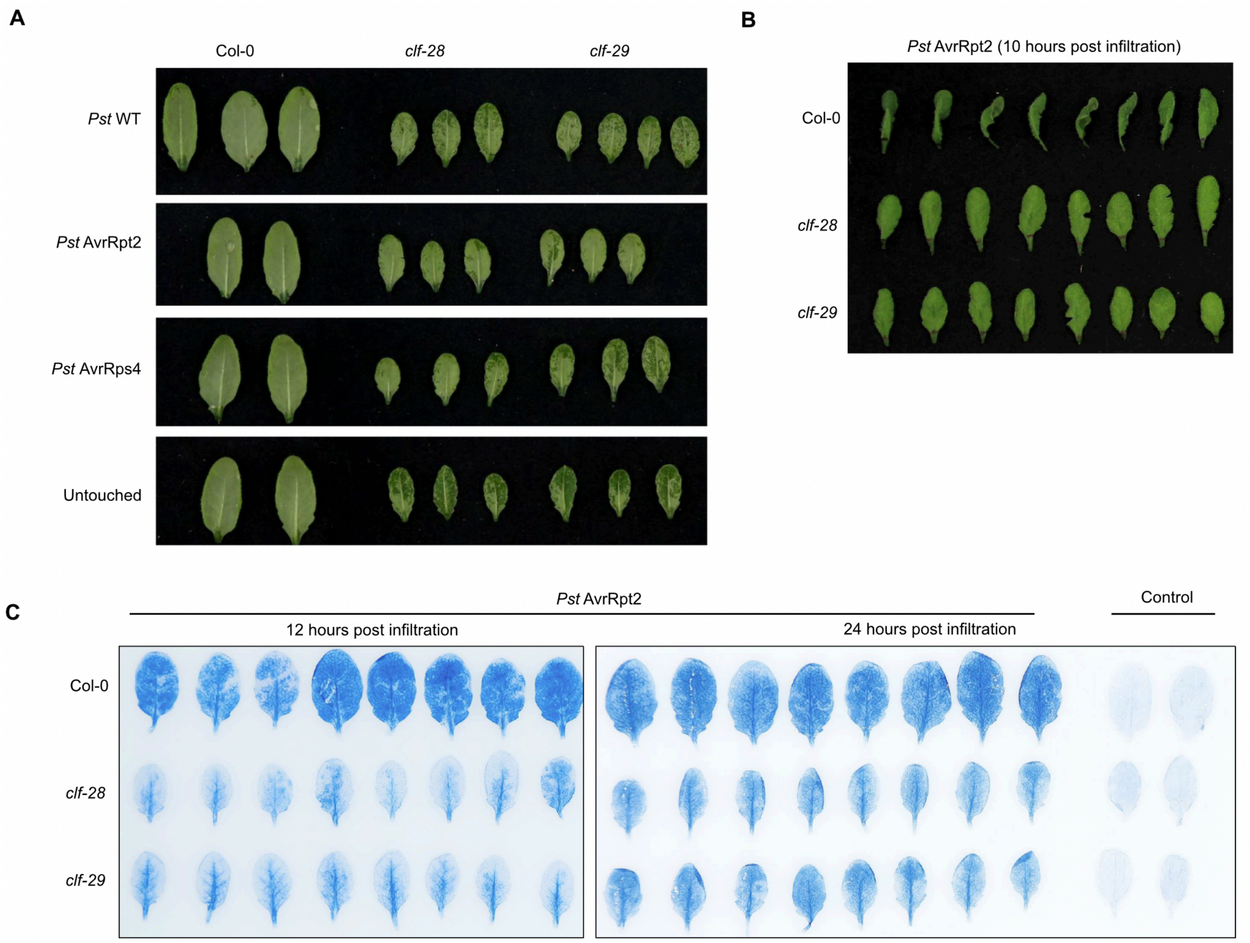


**Figure S8: Aggravated water-soaking in the Arabidopsis *clf* mutants correlates with delayed HR and cell death upon ETI elicitation.**

(**A**) Leaves of four-week-old Arabidopsis WT, *clf-28* or *clf-29* mutant plants that were either untouched or infiltrated with *Pst* WT, AvrRpt2 or AvrRps4 (1 x $10^6$ CFU/mL). Plants were maintained under high humidity (>95% RH) and photos taken 24 hours post-infiltration. (**B**) Reduced tissue collapse (HR) in the Arabidopsis *clf* mutants compared with WT plants during ETI. Arabidopsis WT, *clf-28* or *clf-29* mutant plants were infiltrated with *Pst* AvrRpt2 (1 x $10^8$ CFU/mL) to elicit a strong HR. Plants were kept under low humidity to allow for HR development. Photos were taken 10 hours post infiltration. (**C**) ETI-induced cell death is delayed in the Arabidopsis *clf* mutants. Leaves of four-week-old Arabidopsis WT, *clf-28* or *clf-29* were infiltrated with *Pst* AvrRpt2 (1 x $10^8$ CFU/mL) or water (control) solution and harvested and stained with trypan blue to observe dead cells. Photos were taken at 12 or 24 hours post infiltration.

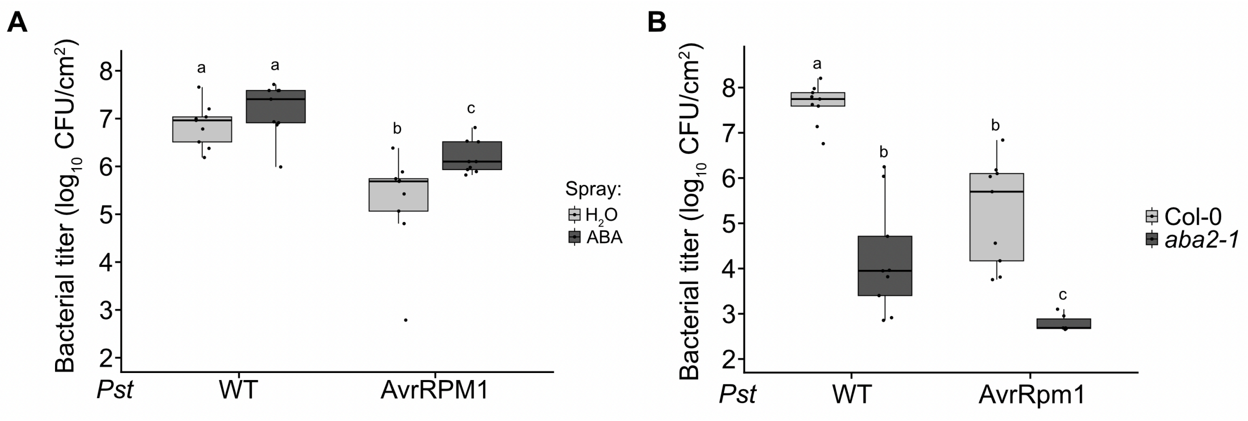


**Figure S9: Impact of ABA on ETI-mediated growth arrest.**

(**A**) Arabidopsis WT leaves were infiltrated with *Pst* DC3000 or AvrRpm1 (1 x $10^5$ CFU/mL). Water or ABA (10 μM) was sprayed on Arabidopsis leaves once per day for the duration of the experiment. Bacterial titer was monitored at 3 dpi. Letters indicate statistical significance ($p < 0.05$) after performing a Pairwise Wilcoxon test followed by a Bonferroni correction. (**B**) Arabidopsis WT or *aba2-1* mutant leaves were infiltrated with *Pst* WT or *Pst* AvrRpm1 (1 x $10^5$ CFU/mL). Bacterial titers were monitored at 3 dpi. Letters indicate statistical significance ($p < 0.05$) after performing a Pairwise Wilcoxon test followed by a Bonferroni correction.

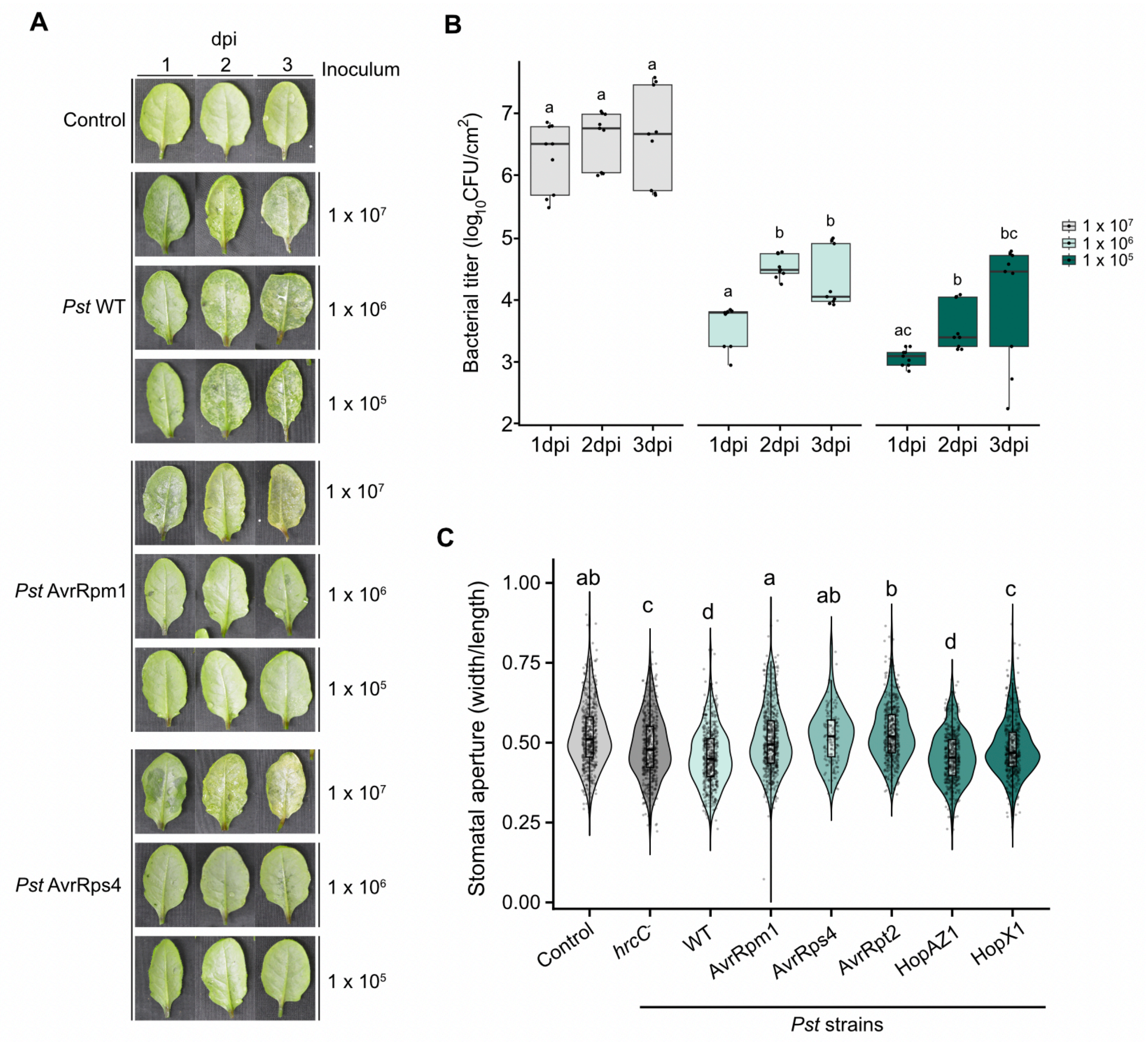


**Figure S10: Influence of bacterial inoculum on water-soaking symptoms, bacterial growth and stomatal aperture.**

(**A**) High inoculum of avirulent *Pst* strain infiltration leads to water-soaking-like symptoms. Arabidopsis WT leaves were infiltrated with a control solution (10 mM $MgCl_2$), *Pst* WT, AvrRpm1 or AvrRps4. Inocula of infiltrated bacteria are indicated in the panel. Photos were taken at either 1, 2 or 3 days post inoculation (dpi). (**B**) Bacterial titer of *Pst* AvrRpm1 after infiltration with different inoculum concentrations ($1 \times 10^7$; $1 \times 10^6$; $1 \times 10^5$ CFU/mL) monitored at 3 different timepoints (1, 2 and 3 dpi). Different letters indicate statistical significance ($p < 0.05$) after performing a Kruskal Wallis test followed by a Pairwise Wilcoxon rank-sum with Bonferroni correction as a post-hoc when Kruskas-Wallis was significant. (**C**) Stomatal aperture of Arabidopsis WT plants infiltrated with control (10 mM $MgCl_2$) or the different *Pst* strain identified in the panel ($1 \times 10^7$ CFU/mL). Stomatal aperture was measured at 1 dpi by scoring their width against their length as obtained as images by epifluorescence microscopy. Different letters indicate statistical significance ($p < 0.05$) after performing a Pairwise Wilcoxon rank-sum (Mann-Whitney U) followed by a Bonferroni correction.

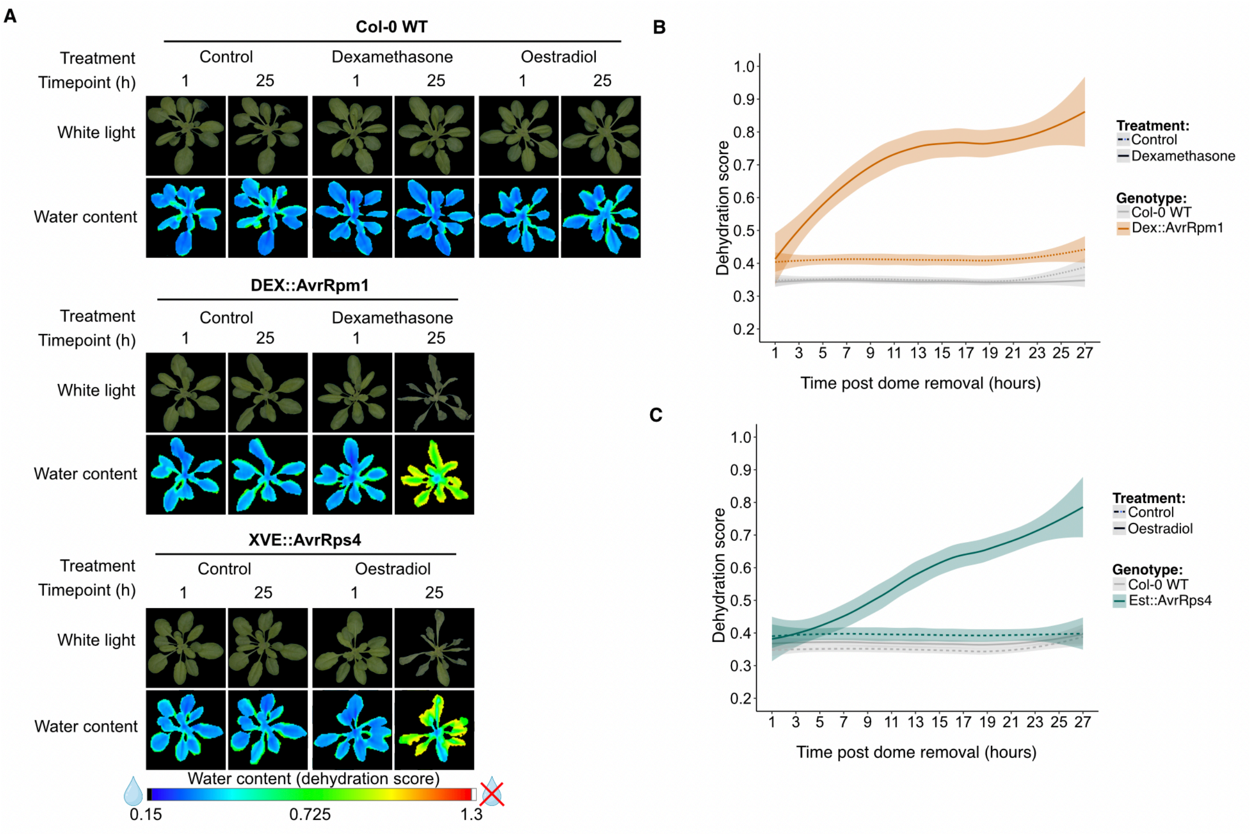


**Figure S11: Plant tissue desiccation in bacteria-free ETI-inducible systems**

(**A**) ETI elicitation post-induction drives tissue desiccation. Photos of Arabidopsis WT or Avr-inducible transgenic plants under white light or hyperspectral imaging of water content at 1 hour or 25 hours post-induction. All plants were sprayed with either water (control), dexamethasone (DEX; 5 µM) for DEX::AvrRpm1 lines or estradiol (XVE; 5 µM) for XVE::AvrRps4 lines. (**B-C**) Dehydration score over time representing water content in Arabidopsis Avr-inducible plants after induction. Dehydration score measured using hyperspectral imaging of water content over 27 hours post-induction as in (**A**).

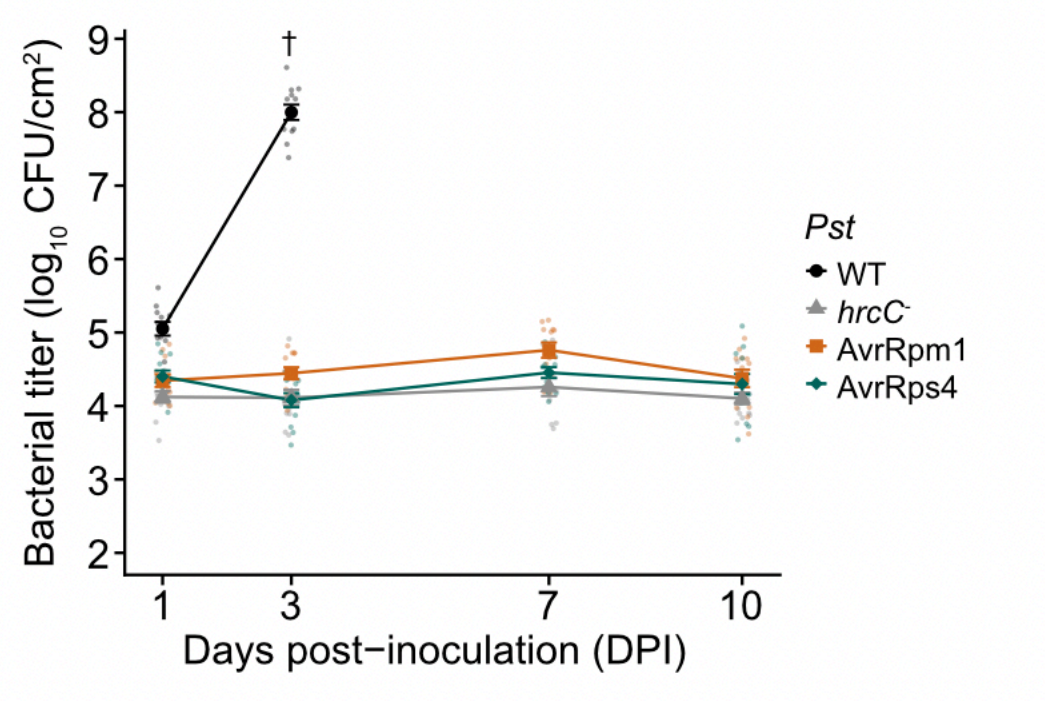


**Figure S12: Population dynamics of virulent, non-virulent and avirulent *Pseudomonas syringae* strains in Arabidopsis leaves.**

Non-virulent and avirulent bacterial strains experience a growth stasis rather than active microbial killing during plant immunity. Arabidopsis WT plants were infiltrated with *Pst* WT, *hrcC*-, AvrRpm1 or AvrRps4 (1 x $10^5$ CFU/mL). Plants were allowed to return to the pre-infiltration state (NF) before being domed to maintain high relative humidity levels. Bacterial titer was evaluated at 1, 3, 7 and 10 days post inoculation.

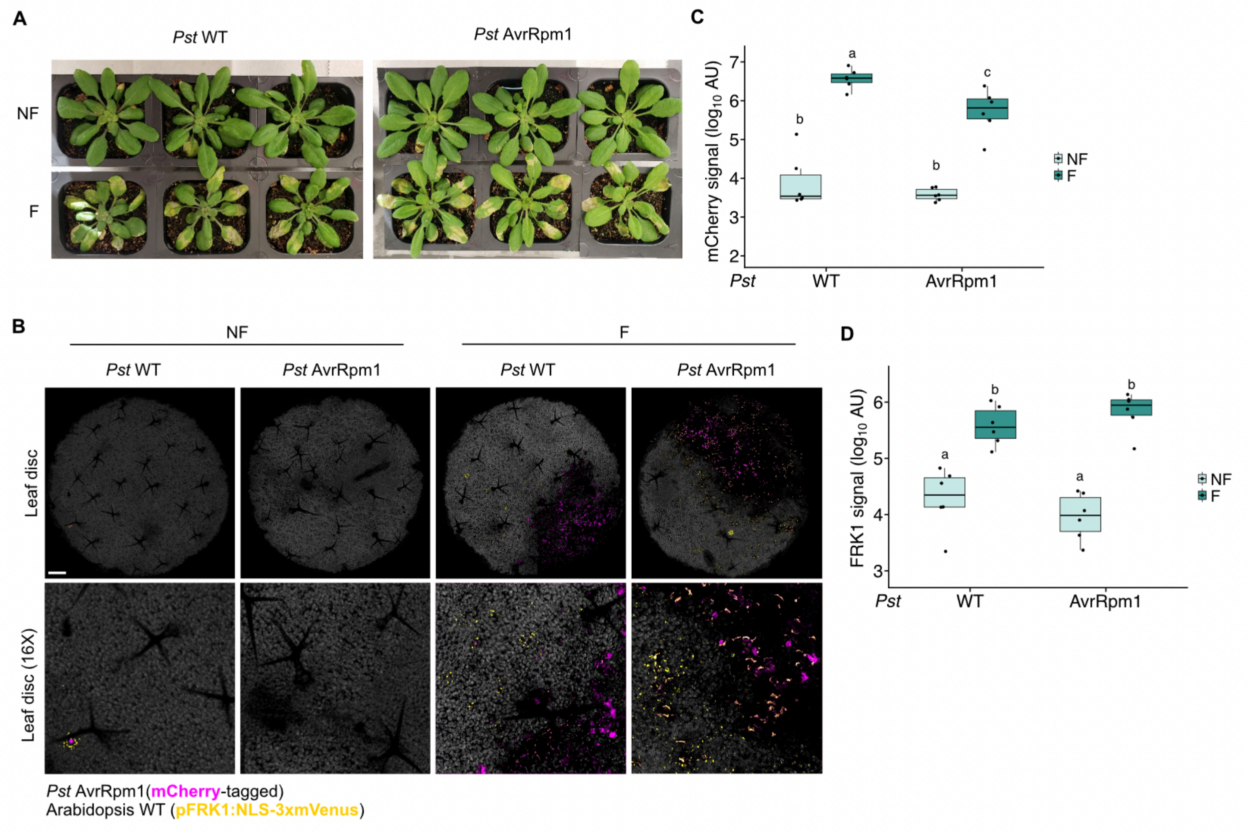


**Figure S13: Loss of disease-resistance in flooded leaves of plants that were surface-inoculated with bacteria.**

(**A**) Four-week-old Arabidopsis WT leaves (carrying the immune marker reporter *pFRK1:NLS-3xmVenus*) were infiltrated with $ddH_2O$. Plants were allowed to evaporate the infiltrated water (NF) or not (F) before being sprayed with mCherry-tagged *Pst* WT or *Pst* AvrRpm1 ($1x10^8$ CFU/mL with added 0.025% Silwet-77). Disease symptoms were then photographed at 3 dpi. (**B**) Confocal microscopy images of Arabidopsis plants carrying the immune transcriptional reporter transgene *pFRK1:NLS-3xmVenus* after surface-inoculation by spraying *Pst* WT or *Pst* AvrRpm1 (mCherry-tagged). (**C**) Average mCherry signal per confocal images, representing bacterial signal as a log10 arbitrary unit (AU). Different letters indicate statistical significance ($p < 0.05$) after performing a Wilcoxon rank-sum (Mann-Whitney U), two-sided with a Bonferroni correction. (**D**) Average *pFRK1:NLS-3xmVenus* signal per confocal images, representing immunity signal as a log10 arbitrary unit (AU). Different letters indicate statistical significance ($p < 0.05$) after performing a Wilcoxon rank-sum (Mann-Whitney U), two-sided with a Bonferroni correction.

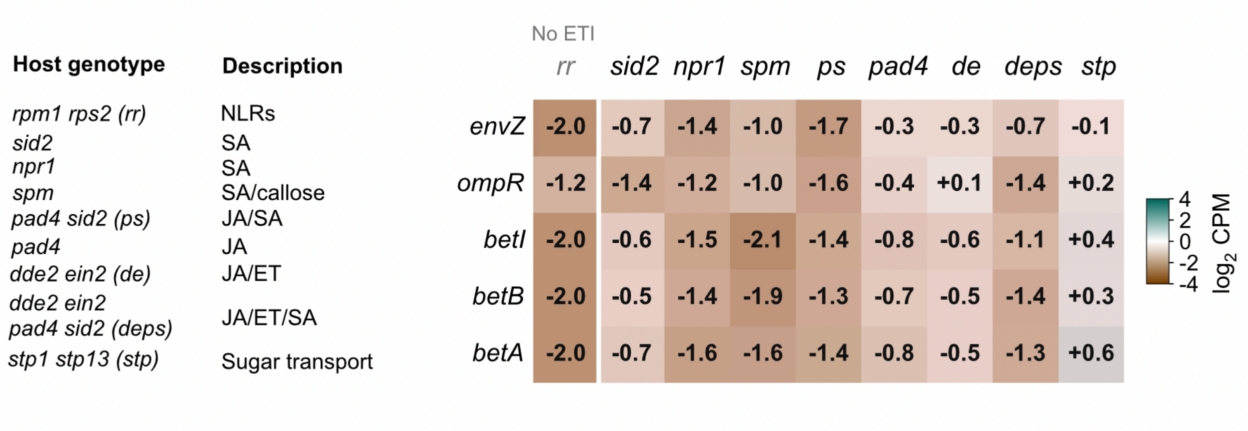


**Figure S14: The ETI-dependent osmotic stress response is dependent on downstream ETI signaling.**

NLR-triggered osmotic stress response in *Pst* AvrRpt2 largely depends on the salicylic acid pathway. Heatmap (Mean log2 count per million) of key selected genes regulated in *Pst* AvrRpt2, compared to *Pst* WT, associated with bacterial osmotic stress response in Arabidopsis mutants for immune perception, phytohormones biosynthesis, callose deposition and/or sugar transport. Data were re-analyzed from a previously published work (Nobori et *al*., 2018).
